# *Bergeyella cardium variant* induces unique cytoplasmic vacuolization cell death

**DOI:** 10.1101/2024.09.24.614850

**Authors:** Rudi Mao, Hongwei Pan, Luyu Yang, Zhen Li, Xinran Yu, Yanfeng Li, Ying Chen, Yang Yu, Wei Wang, Chengjiang Gao, Jun Peng, Tao Xu, Yi Zhang, Xiaopeng Qi

**Author notes:** Correspondence to: Tao Xu, Yi Zhang, Xiaopeng Qi. Co-first author (Rudi Mao, Hongwei Pan).

## Abstract

Bacterial pathogens have evolved multiple mechanisms to modulate host cell death, evade host immunity, and establish persistent infection. Here, we show that an emerging pathogen *Bergeyella cardium variant* (*BCV*) survives serum killing, inducing unique cytoplasmic vacuolization cell death in the majority of cells and apoptosis in a small number of bystanders. The cytoplasmic vacuolization cell death triggered by *BCV* is characterized by fused lysosome-associated termination and is inhibited by the sodium channel inhibitor amiloride. Endosomal solute carrier family 9 member A9 (SLC9A9) was identified as a critical regulator in the process of *BCV*-triggered cytoplasmic vacuolization cell death. Moreover, outer membrane vesicles (OMVs) or transfection of barrel-like membrane proteins lipocalin, β-barrel, and PorV dramatically induced cytoplasmic vacuolization. *BCV* hijacked SLC9A9 by maintaining its high expression, increasing cytoplasmic vacuolization, and enhancing pathogenicity. These findings contribute to the development of novel approaches to modulate cytoplasmic vacuolization cell death and treat infectious diseases.

**Highlights:**

1. *BCV* triggers unique cytoplasmic vacuolization cell death.
2. SLC9A9 is essential for *BCV*-induced cytoplasmic vacuolization cell death.
3. OMVs and the transfection of barrel-like proteins derived from *BCV* induce cytoplasmic vacuolization cell death.
4. *BCV* increases pathogenicity via cytoplasmic vacuolization cell death.

## Introduction

Host cell death is an intrinsic immune response that plays dual roles in host defence against bacterial infection^1^. Bacterial pathogens have evolved distinct strategies to manipulate host cell death and survival pathways, creating an intracellular replication niche and promoting dissemination^2^. The major programmed cell death pathways, including apoptosis, necroptosis, and pyroptosis have been extensively studied, and many virulence factors of bacterial pathogens can act through these pathways to increase pathogenicity^3–5^. Cytoplasmic vacuolization is a morphological phenomenon characterized by the dilation and fusion of the endoplasmic reticulum or endosomal–lysosomal organelles, which can be transient or irreversible depending on the properties of chemical and infectious agents^6^. However, the relationship between vacuolization and cell death, particularly whether the vacuolization phenomenon is a programmed cell death process, remains unknown. In addition, the role of cytoplasmic vacuolization in the inflammatory response and immune evasion has not been investigated.

We previously isolated an emerging pathogen strain with a flat dry and rough morphotype colony, *Bergeyella cardium* (*BC*), from a patient with infectious endocarditis^7^. In the present study, we observed that a variant strain of *BC* (*BCV*) had a smooth morphotype and increased resistance to serum complement-dependent clearance. Whole-genome sequencing revealed high sequence identity between *BC* and *BCV*. Notably, *BCV* infection triggered robust cytoplasmic vacuolization cell death and minor apoptosis in different cells, whereas *BC* infection activated necroptosis. The vacuoles induced by *BCV* were colocalized with the late endosomal and lysosomal markers Rab7 and LAMP1. Furthermore, *BCV*-triggered cytoplasmic vacuolation cell death was dependent on SLC9A9. Both OMVs derived from *BCV* and transfection of lipocalin, β-barrel, and PorV of *BCV* induced the formation of cytoplasmic vacuolization. The inflammatory responses triggered by *BCV* were much weaker than those triggered by *BC*. In contrast, the intracellular replication and in vivo pathogenicity of *BCV* were significantly stronger than those of *BC*. Our work offers novel insights into the mechanisms of cytoplasmic vacuolization cell death and provides potential therapeutic targets for the treatment of infectious diseases.

## Results

### *BCV* is more resistant to serum killing than *BC* and is prone towards high pathogenicity

To investigate the detailed bacterial characteristics and pathogenesis of *BC*, which was previously isolated from a patient with infectious endocarditis^7^, we spread the bacterial stock on Columbia blood agar plates. Interestingly, we identified two distinct colony morphologies (Figure 1A). The *BC* colonies were flat, dry, and rough, whereas the *BCV* colonies were raised and smooth (Figure 1A). We performed whole-genome sequencing of *BC* and *BCV* colonies, and the circular genome lengths of *BC* and *BCV* were 2,036,890 bp and 2,036,968 bp, respectively (Figure 1B; GenBank CP029149 and CP114055). The genomic homology between *BC* and *BCV* was approximately 99.99%. We compared the *BC* and *BCV* genomes and confirmed that the *BCV* genome harbored three insertions located in the noncoding region, the *PorV* gene and a hypothetical gene of the *BC* genome (Figures S1A-S1F). The 24–bp insertion in the *PorV* gene caused an 8-amino-acid insertion in the PorV protein of *BCV*, resulting in a PorV protein longer than that of *BC* (Figures S1C and S1D). The 13–bp insertion in the hypothetical gene of *BC* caused a frameshift mutation, which produced a new BatD family protein in *BCV* (Figures S1E and S1F). To characterize the phenotypes of *BC* and *BCV* in detail, we performed electron microscopy analysis. Scanning electron microscopy (SEM) and transmission electron microscopy (TEM) revealed that *BCV* bacteria had multiple membrane puff-ups and blebbing (Figure 1C). Serum resistance of bacteria is essential for their persistence in the bloodstream to cause infection and for the development of antibiotic tolerance, which is recognized as a stepping stone towards antibiotic resistance^8–10^. Notably, the capacity for serum resistance in *BCV* was much greater than that in *BC* (Figures 1D and S2A), whereas antibiotic resistance was comparable between *BC* and *BCV* (Figure S2B).

**Figure 1.**
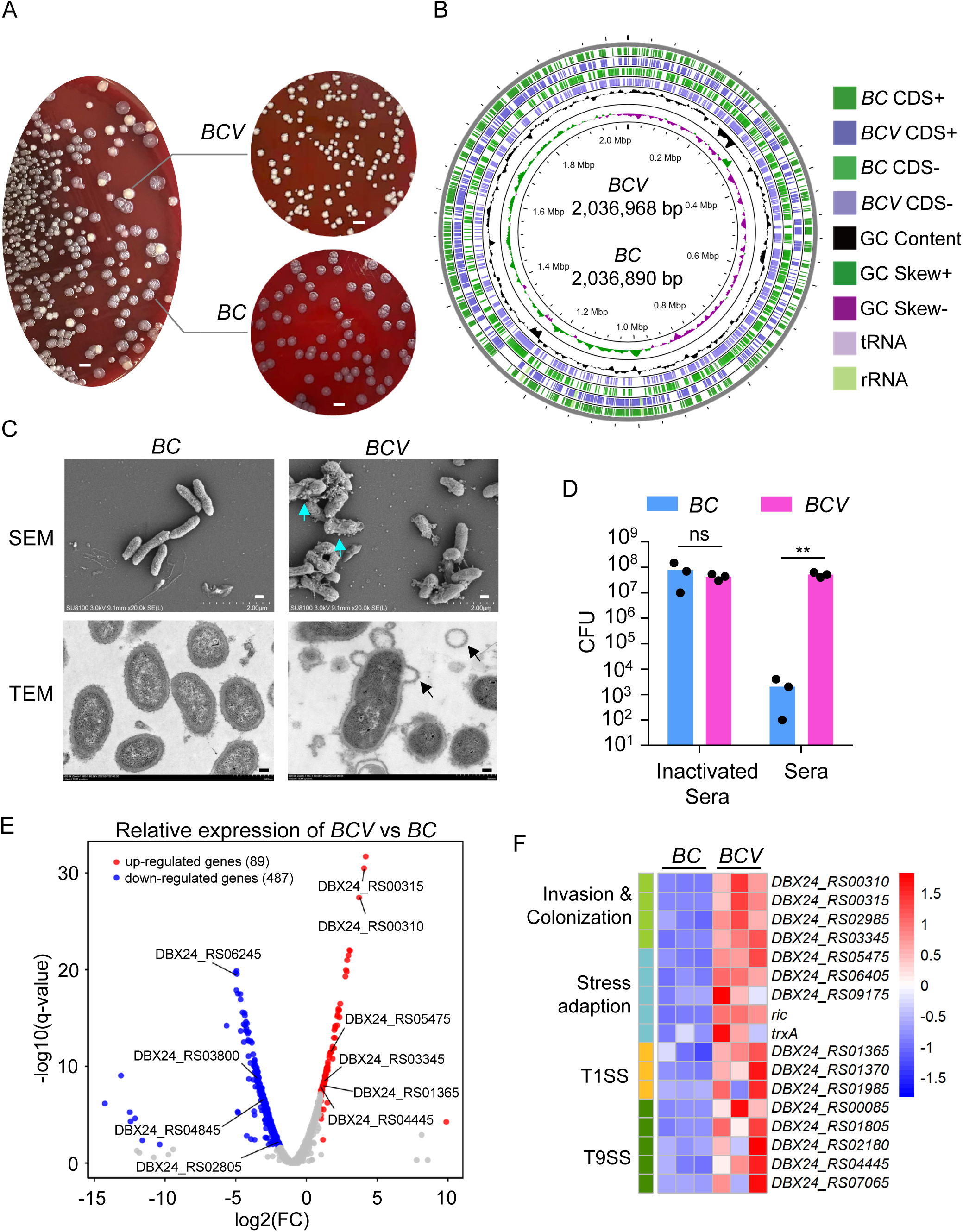
Characterization of *BC* and *BCV*. **A**, Bacterial colony morphology of *BC* and *BCV*. *BC* and *BCV* were cultured on Columbia blood agar plates for 96 hours. Scale bars, 3 mm. **B**, Circular genome map of *BCV* with its alignment with *BC*. High-quality and scalable circular genome maps were generated using the Java package CGView to display the alignment of the *BCV* genome with the *BC* genome. The circular genome map includes the following, from outer to inner rings: *BC* CDS+, coding sequences (CDS) on the forward strand of the *BC* genome; *BCV* CDS+, CDS on the forward strand of the *BCV* genome; *BC* CDS-, CDS on the reverse strand of the *BC* genome; *BCV* CDS-, CDS on the reverse strand of the *BCV* genome; GC content, which is useful for identifying genomic islands and horizontal gene transfer events; GC skew, indicating the over- or under-abundance of G or C between the leading and lagging DNA strands, which is often used to identify the origin and terminus of replication; and the locations of tRNAs and rRNAs in the genome. The *BC* genome is 2,036,890 bp long, and the *BCV* genome is 2,036,968 bp long. **C**, SEM and TEM analysis of *BC* and *BCV*. *BC* and *BCV* were cultured on Columbia blood agar plates for 96 hours, followed by culture in BACTEC™ Lytic Anaerobic media for 16 hours. Equal numbers of *BC* and *BCV* were fixed and sectioned for SEM and TEM analysis. The arrows indicate puff-up and blebbing of *BCV*. Scale bars, 2 μm for SEM and 1 μm for TEM. **D**, Colony formation analysis of *BC* and *BCV* treated with human sera. Sera were inactivated at 56 °C for 30 min. *BC* and *BCV* were cultured on Columbia blood agar plates for 96 hours, followed by growth in BACTEC™ Lytic Anaerobic media for 16 hours, and equal numbers of *BC* and *BCV* were incubated with normal and heat-inactivated sera for 2 hours at 37 °C. Sera-treated *BC* and *BCV* were grown on Columbia blood agar plates for 96 hours, and the numbers of bacteria were enumerated. Each dot represents an individual experiment (n=3 biologically independent samples). **E**, RNA-seq analysis of gene expression in *BC* and *BCV*. *BC* and *BCV* were cultured on Columbia blood agar plates for 96 hours, after which RNA-seq analysis was performed. Volcano plot showing the distribution of upregulated (red) and downregulated (blue) genes at the transcriptional level in *BCV* compared with *BC*. A generalized linear model (GLM) framework was utilized to perform the differential expression analysis, including the log2-fold change (log2FC), p value and q value for each gene. The q value is the p value adjusted for the false discovery rate (FDR). **F**, Heatmap analysis of genes in (**E**) that were highly expressed in *BCV* but not in *BC*. Data are from 3 independent experiments (**D**-**F**) or representative of 3 independent experiments with similar results (**A**, **C**). Data represent Mean ± SEM for (**D**), **P < 0.01, by 2-sided Student’s t test without multiple-comparisons correction.

To define the genes differentially expressed between *BC* and *BCV*, we performed RNA sequencing analysis of *BC* and *BCV* cultured on Columbia blood agar plates for 96 hours. Overall, 89 genes and 487 genes were upregulated and downregulated, respectively, in *BCV* (Figure 1E and Table S1). KEGG enrichment analysis revealed that metabolism pathways, including the pentose phosphate, TCA cycle, and methane pathways, were upregulated in *BCV*, whereas biosynthesis pathways, including ribosome and aminoacyl-tRNA pathways, were upregulated in *BC* (Figure S2C). Notably, the genes involved in bacterial invasion and colonization, stress adaptation, and secretion systems, including types I and IX, were highly expressed in *BCV* (Figures 1F and S2F). Conversely, the genes encoding proteins involved in immunogenicity and pathogenicity, such as lipopolysaccharide (LPS) and lipoteichoic acid (LTA) biosynthesis, were dramatically downregulated in *BCV* (Figures S2D and S2E). PorV is a shuttle protein in the outer membrane of the type IX secretion system (T9SS) that mediates the secretion of major virulence factors^11^. To investigate whether the insertion of 8 amino acids in PorV of *BCV* (residues 344–351) led to its structural difference from *BC* (Figure S3A), we performed structure prediction analysis using trRosseta^12^. According to the predicted structure, *BCV* PorV folded into a 14-strand β-barrel, and residues 344–351 folded into the β15 of the β-barrel (denoted ‘*BCV* β15’). *BC* PorV was also predicted to be a 14-strand β-barrel. However, since the residues corresponding to ‘*BCV* β15’ were missing, residues at the N- and C-termini of ‘*BCV* β15’ were forced into *BC* β15 in the predicted structure (Figures S3B and S3C), which altered the residue composition of the β15 strand. Notably, the predicted structure of *BC* PorV was less reliable in the β14–β15 region, as shown by the lower LDDT in this region (Figures S3B and S3C). Therefore, compared with *BCV* PorV, deletion of ‘*BCV* β15’ in *BC* PorV caused structural changes in PorV, which might affect the secretion of virulence factors through interactions with binding partners. Overall, these results suggested that *BCV* could survive in the human bloodstream and might represent a more virulent *BC* variant strain.

### *BCV* induces lysosomal fusion-mediated cytoplasmic vacuolization cell death

To investigate the host cell death response triggered by *BC* and *BCV* infection, we treated bone marrow-derived macrophages (BMDMs) with *BC* or *BCV* for different durations. Notably, *BCV* infection triggered cytoplasmic vacuolization beginning at 6 hours post infection, significant cytoplasmic vacuolization at 10 hours post infection, and large vacuoles and single vacuole-occupied cells at 20 and 40 hours post infection (Figures 2A and S4A). However, this phenomenon was not observed in *BC*-infected BMDMs (Figure 2A). In addition to BMDMs, *BCV* infection also triggered cytoplasmic vacuolation in L929 cells, immortalized BMDMs (iBMDMs), and the human cardiomyocyte cell line AC16 (Figures S4B-S4F). To closely examine vacuolization, we performed TEM analysis of *BC*- and *BCV*-infected BMDMs. Notably, *BCV* infection caused the formation of multiple single membrane-bound vacuoles at 10 hours post infection (Figure 2B). The size of the vacuoles was enlarged, and the number was reduced due to the fusion of vacuoles during the later phase of infection (Figures 2B and S4G). The process of vacuole fusion and enlargement was also confirmed by live-cell imaging analysis (Figure 2C). The vacuoles triggered by *BCV* were labelled with LAMP1 and Rab7 but not the early endosome markers Rab5 or EEA1 (Figures 2D and S5A-S5C), which indicated that the vacuoles were derived from lysosomes and late endosomes. However, necroptotic cell death characterized by robust membrane rupture was observed in *BC*-infected BMDMs at later time points (Figure 2B). Remarkably, the cell death caused by *BCV* infection was much more severe than that caused by *BC* infection, as determined by real-time quantification of cell death and an LDH release assay (Figures 2E-2G). In addition, the density of *BCV* infection-induced vacuole-containing cells was lower than that of normal BMDMs, and vacuolated cells were separated at the middle layer of the 40% Percoll gradient (Figure S5D). Thus, we termed this type of cell death Fused LysosOme-Associated Termination (floatptosis).

**Figure 2.**
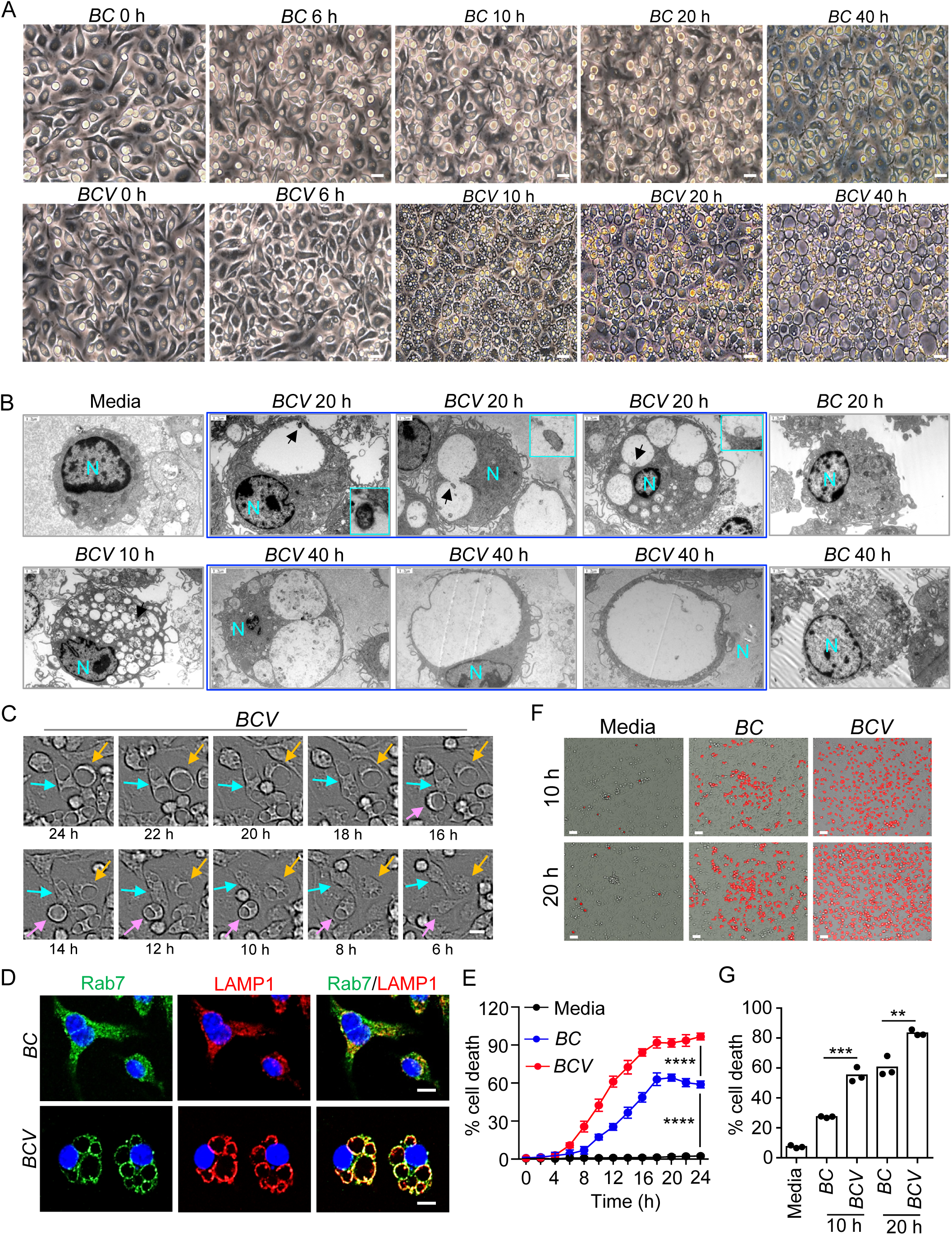
*BCV* infection triggers cytoplasmic vacuolization cell death. **A**, Microscopic analysis of WT BMDMs infected with *BC* or *BCV* (400 MOI) for the indicated times. Scale bars, 30 μm. **B**, TEM analysis of WT BMDMs infected with *BC* or *BCV* (400 MOI) for the indicated times. N indicates the cell nucleus, arrows indicate the intracellular *BCV*, and enlarged images are shown. Scale bars, 1.2 μm. **C**, Live-cell imaging of WT BMDMs infected with *BCV* (400 MOI) for the indicated times. Scale bar, 20 μm. **D**, Confocal microscopy analysis of Rab7 and LAMP1 in *BC*- and *BCV*-infected (400 MOI) WT BMDMs for 20 hours. Scale bars, 20 μm. **E**, Real-time quantitative live-cell imaging and analysis of cell death in uninfected WT BMDMs and WT BMDMs infected with *BC* or *BCV* (400 MOI) (n=16 random fields; 3 independent experiments). **F**, Representative images of PI staining of uninfected BMDMs and BMDMs infected with *BC* or *BCV* (400 MOI) for the indicated times in (**E**). Scale bars, 20 μm. **G**, LDH analysis of WT BMDMs infected with *BC* or *BCV* (400 MOI) for the indicated times (n=3 biologically independent samples). Data are from 3 independent experiments (**G**) or representative of 3 independent experiments with similar results (**A**-**F**). Data represent Mean ± SEM for (**G**), 2-sided Student’s t test without multiple-comparisons correction, two-way ANOVA for (**E**). **P < 0.01, ***P < 0.001, ****P < 0.0001.

### *BCV*-triggered floatptosis is a unique cell death pathway

Type I IFN (IFN-I) signalling and inflammasome activation play important roles in many bacterial pathogen infection-triggered cell death pathways, such as pyroptosis and PANoptosis^4,13,14^. To determine whether IFN-I signalling and inflammasome activation were involved in the *BCV*-triggered floatptosis pathway, WT, *Ifnar^-/-^, Asc^-/-^, and Aim2^-/-^Nlrp3^-/-^* BMDMs were infected with *BC* or *BCV* (Figure S6A). The results showed that the levels of cytoplasmic vacuolization and LDH release induced by *BCV* infection were comparable among these samples (Figures S6A-S6C), indicating that IFN-I and inflammasome activation were dispensable for *BCV*-triggered floatptosis. To further examine whether this type of cytoplasmic vacuolization was associated with apoptosis or other types of cell death, we analysed the hallmarks of different types of cell death via protein expression and activation analyses of caspase-3 for apoptosis, caspase-1 for pyroptosis, and phosphorylation of MLKL and RIP3 for necroptosis in *BC*- and *BCV*-infected BMDMs. Indeed, *BCV* infection also triggered caspase-3 activation at later time points but did not affect necroptosis or pyroptosis (Figures 3A, S6D and S6E). *BC* infection induced the activation of necroptosis via the phosphorylation of MLKL and RIP3, which was consistent with the TEM analysis of *BC*-infected BMDMs (Figures 2B and S6D). Notably, TEM analysis of *BCV*-infected BMDMs at later time points revealed that *BCV* infection activated two types of cell death in different cells: major floatptosis (78.65%) and minor apoptosis (21.35%) (Figure 3B). Approximately 20% of the apoptosis induced by *BCV* but not *BC* in BMDMs was confirmed by Annexin V staining (Figure S6F). Furthermore, the supernatant without viable bacteria from *BCV*-but not *BC*-infected BMDMs induced caspase-3 activation (Figure S6G), whereas the supernatant from either *BCV*- or *BC*-infected BMDMs did not induce obvious cytoplasmic vacuolization (Figures S6H and S6I). Thus, we performed mass spectrometry (MS) analysis of supernatants from *BCV*- and *BC*-infected BMDMs and found that lysosomes and lysosome-associated proteins, such as multiple cathepsin proteins, were enriched in the supernatants of *BCV*-treated BMDMs compared with *BC*-treated cells (Figure S6J and Table S2), suggesting that lysosomes and lysosome-associated proteinases enriched in the supernatant of *BCV*-infected BMDMs might contribute to the apoptosis of bystander cells.

**Figure 3.**
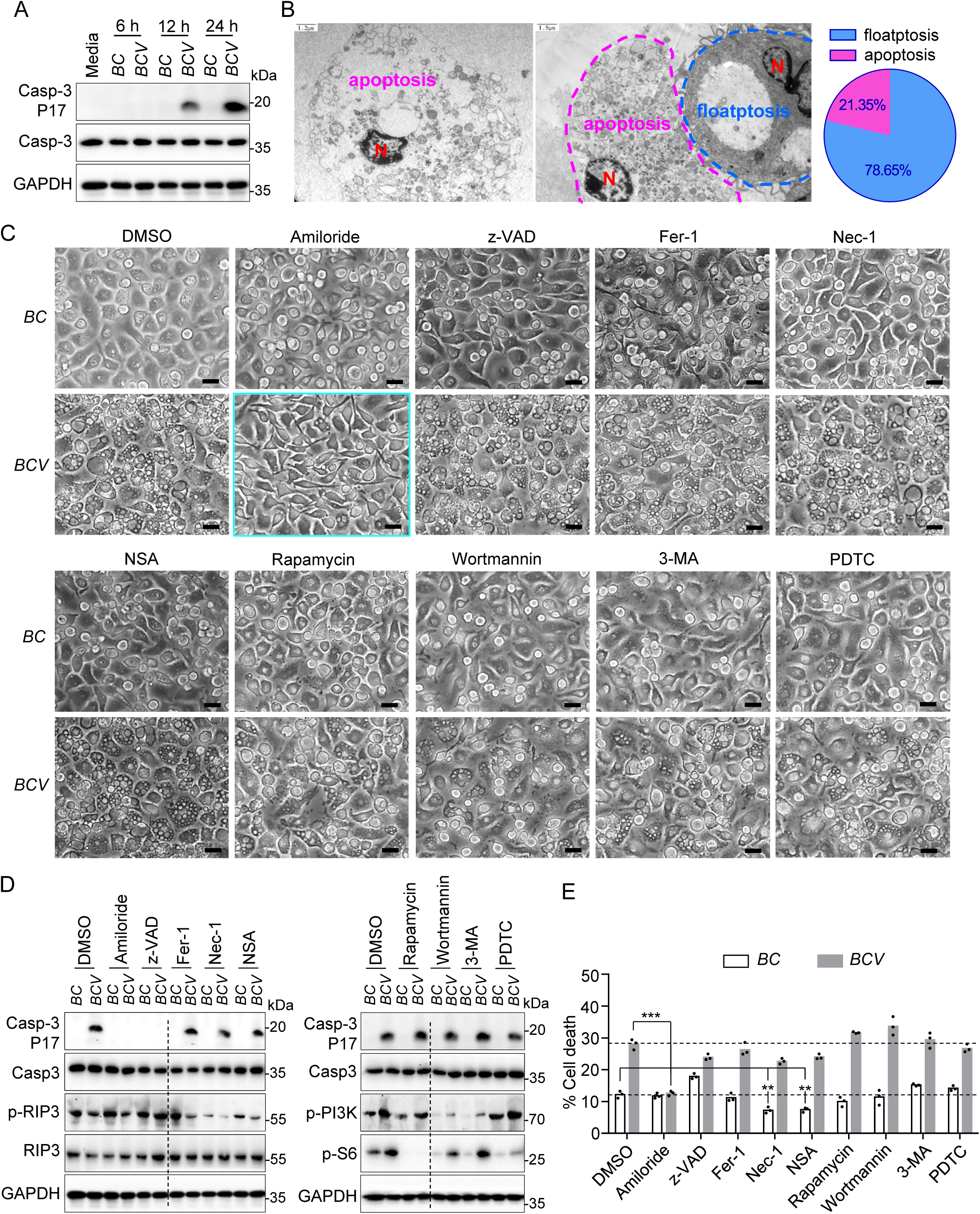
Amiloride inhibits *BCV*-triggered cytoplasmic vacuolization cell death. **A**, Immunoblot analysis of caspase-3 and cleaved caspase-3 (P17) in WT BMDMs infected with *BC* or *BCV* (400 MOI) for the indicated times. **B**, TEM analysis of WT BMDMs infected with *BCV* (400 MOI) for 30 hours (left and middle) and quantification of the percentage of floatptosis and apoptosis within the dead cell population (right). N indicates the cell nucleus. A total of 89 dead cells were analysed. Scale bars, 1.2 μm for the left and 1.5 μm for the middle. **C**, Microscopic analysis of WT BMDMs infected with *BC* or *BCV* (400 MOI) in the presence of amiloride hydrochloride (Amiloride, 0.5 μM), z-VAD (25 μM), ferrostatin-1 (Fer-1, 1 mM), necrostatin-1 (Nec-1, 100 μM), necrosulfonamide (NSA, 500 nM), Rapamycin (500 nM), Wortmannin (0.2 μM), 3-Methyladenine (3-MA, 5 mM), and pyrrolidinedithiocarbamate ammonium (PDTC, 1 μM) for 12 hours. Scale bars, 20 μm. **D**, Immunoblot analysis of caspase-3, cleaved caspase-3 (P17), p-RIP3, RIP3, p-PI3K, and p-S6 in WT BMDMs infected with *BC* or *BCV* (400 MOI) together with various inhibitors in (**c**) for 12 hours. **E**, LDH analysis of WT BMDMs infected with *BC* or *BCV* (400 MOI) in the presence of the indicated inhibitors for 12 hours (n=3 biologically independent samples). Data are from 3 independent experiments (**E**) or representative of 3 independent experiments with similar results (**A**-**D**). Data represent Mean ± SEM for (**E**), 2-sided Student’s t test without multiple-comparisons correction, **P < 0.01, ***P < 0.001.

To further examine whether *BCV*-triggered cytoplasmic vacuolization was dependent on other cell death pathways and intracellular events, WT BMDMs were infected with *BC* or *BCV* in the presence or absence of different types of inhibitors, including the pancaspase inhibitor z-VAD for apoptosis inhibition, the RIPK1 inhibitor necrostatin-1 (Nec-1) and MLKL inhibitor necrosulfonamide (NSA) for necroptosis inhibition, ferrostatin-1 (Fer-1) for ferroptosis inhibition, pyrrolidinedithiocarbamate ammonium (PDTC) for NF-κb inhibition, 3-methyladenine (3-MA) and wortmannin for PI3K and autophagy inhibition, rapamycin for mTORC1 inhibition and autophagy activation, and amiloride hydrochloride for sodium channel inhibition. Notably, amiloride dramatically inhibited *BCV*-triggered cytoplasmic vacuolization, caspase-3 activation, and LDH release (Figures 3C-3E and S7A). Conversely, z-VAD inhibited caspase-3 activation but not cytoplasmic vacuolization or cell death (Figures 3C-3E and S7A), indicating that *BCV*-triggered cytoplasmic vacuolization cell death was not dependent on apoptosis. In addition, inhibitors of necroptosis, ferroptosis, autophagy, mTORC1, or NF-κb did not affect *BCV* infection-triggered cytoplasmic vacuolization, caspase-3 activation or LDH release (Figures 3C-3E and S7A). The necroptosis inhibitors Nec-1 and NSA clearly inhibited *BC* infection-triggered cell death (Figures 3D and 3E). Overall, these results indicated that *BCV* infection induced two separate cell death pathways: floatptosis and, to a minor extent, apoptosis. It is possible that apoptosis may be a downstream secondary effect of floatptosis induced by *BCV*, and ion channels are important for *BCV*-triggered floatptosis.

### SLC9A9 is essential for *BCV*-induced floatptosis

To explore the mechanism underlying *BCV* infection-induced floatptosis, we performed a genome-wide transcriptional analysis of *BC*- and *BCV*-infected BMDMs. *BCV* infection triggered the enrichment of genes associated with endosomes, lysosomes, and metabolism (Figures 4A, S7B, and Table S3). However, the expression levels of genes involved in inflammatory response pathways, including TNF, chemokine, NF-κB, and MAPK signalling, were dramatically reduced in *BCV*-compared with *BC*-infected BMDMs (Figures S7C-S7E and Table S3). Given that inhibitors targeting ion channels have suppressive effects on the process of *BCV* infection-induced cytoplasmic vacuolization, we found that certain genes encoding membrane proteins and transporters that were localized to intracellular compartments, especially solute carriers (SLCs), were highly expressed in BMDMs infected with *BCV* but not *BC* (Figures 4B and S7F).

**Figure 4.**
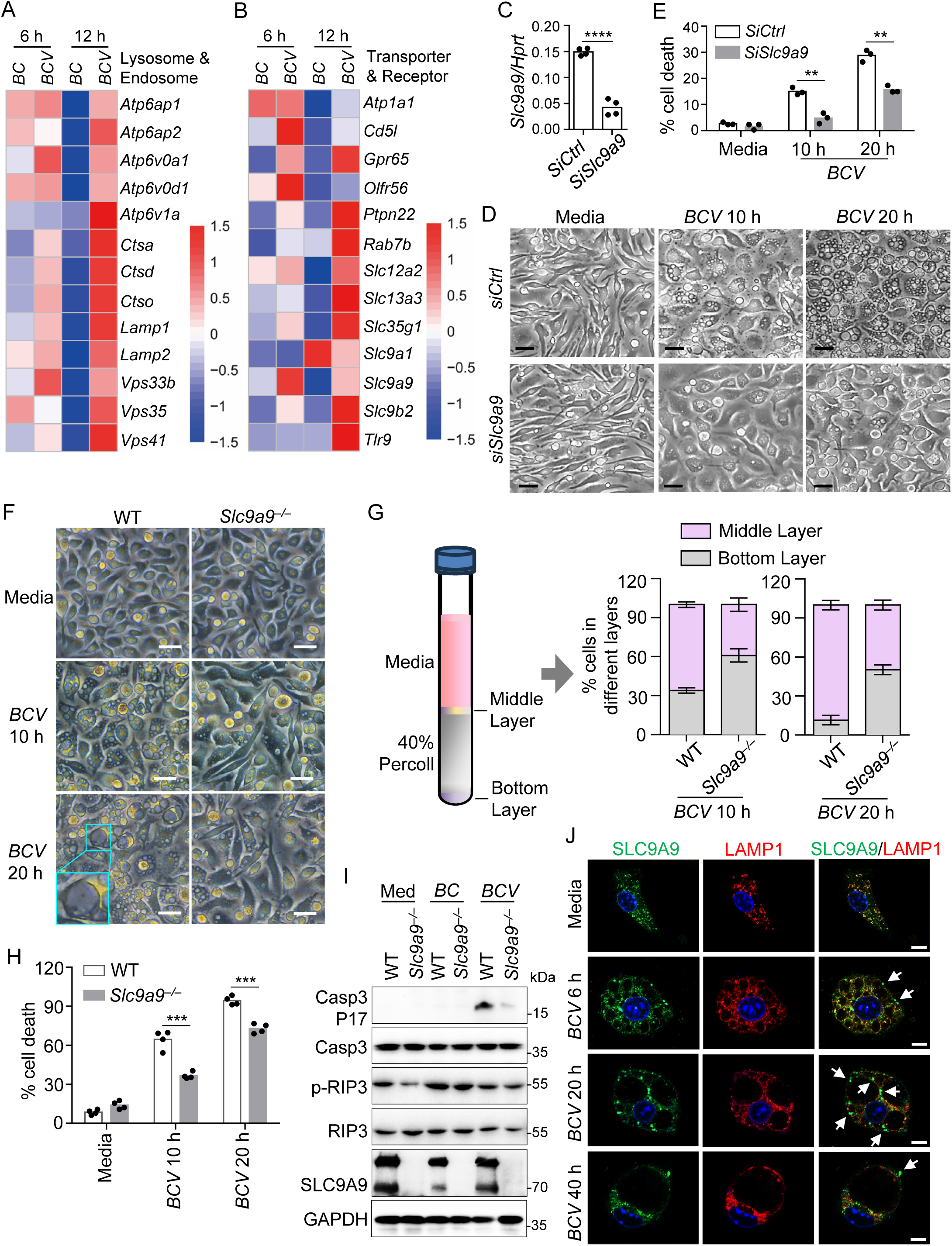
SLC9A9 mediates *BCV* infection-triggered cytoplasmic vacuolization cell death. **A**, **B**, RNA-seq analysis of the gene expression of WT BMDMs infected with *BC* or *BCV* (400 MOI) for 6 and 12 hours. Heatmap analysis of endosome- and lysosome-associated genes (**A**) and genes encoding receptors and transporters (**B**) with increased expression in *BCV*-infected BMDMs. **C**, Quantitative RT-PCR analysis of *Slc9a9* expression in *siCtrl*- and *siSlc9a9*-transfected BMDMs (20 μM siRNA for each) for 20 hours (n=4 technical replicates; 3 independent experiments). **D**, Microscopic analysis of siRNA-knockdown BMDMs (*siCtrl* and *siSlc9a9*) infected with *BCV* (400 MOI) for the indicated times. Scale bars, 30 μm. **E**, LDH analysis of *siCtrl*- and *siSlc9a9*-transfected BMDMs in (**D**) (n=3 biologically independent samples). **F**, Microscopic analysis of WT and *Slc9a9^-/-^* BMDMs infected with *BCV* (400 MOI) for the indicated times. Scale bars, 30 μm. **G**, Quantification of the level of cytoplasmic vacuolization in *BCV*-infected WT and *Slc9a9^-/-^* BMDMs (400 MOI) by 40% Percoll gradient separation (n=3 biologically independent samples). **H**, LDH analysis of *BCV*-infected WT and *Slc9a9^-/-^* BMDMs (400 MOI) for the indicated times (n=4 biologically independent samples). **I**, Immunoblot analysis of caspase-3, cleaved caspase-3 (P17), p-RIP3, RIP3, and SLC9A9 in WT and *Slc9a9^-/-^*BMDMs infected with *BC* or *BCV* (400 MOI) for 12 hours. **J**, Confocal microscopy analysis of SLC9A9 and LAMP1 in *BCV*-infected (400 MOI) WT BMDMs at the indicated times. The arrows indicate SLC9A9 subcellular localization at membrane fusion sites. Scale bars, 10 μm. Data are from 3 independent experiments (**E**, **G**, **H**) or representative of 3 independent experiments with similar results (**C**, **D**, **F**, **I**, **J**). Data represent Mean ± SEM for (**C**, **E**, **G**, **H**), 2-sided Student’s t test without multiple-comparisons correction, **P < 0.01, ***P < 0.001, ****P < 0.0001.

We performed siRNA transfection for 13 candidate genes that were more highly expressed in *BCV*-infected than in *BC*-infected BMDMs (Figures 4B, 4C and S8A). Notably, the cytoplasmic vacuolization and LDH release induced by *BCV* infection were markedly inhibited in *SiSlc9a9*-transfected BMDMs but not in *SiSlc9a1*-, *SiSlc9b2*-, *SiSlc12a2*-, *SiSlc13a3*-, *SiSlc35g1*-, *SiAtp1a1*-, *SiRab7b*-, *SiGpr65*-, *SiCd5l*-, *SiOlfr56*-, *SiPtpn22*-, or *SiTlr9*-transfected BMDMs (Figures 4D, 4E and S8B-S8D). Consistently, the overexpression of *Slc9a9* markedly increased *BCV*-triggered floatptosis by increasing the formation rate and size of vacuoles (Figures S9A-S9D). The knockout of *Slc9a9* in iBMDMs resulted in reduced floatptosis upon *BCV* infection (Figures S9E-S9H).

To provide genetic evidence for the role of SLC9A9 in the process of *BCV*-triggered floatptosis, we generated *Slc9a9^-/-^* mice (Figures S9I-S9K), which were viable with characteristics similar to those of WT mice. SLC9A9 deficiency in BMDMs significantly reduced the degree of *BCV*-triggered floatptosis based on microscopy, quantification by 40% Percoll gradient separation, and LDH release assay analyses (Figures 4F-4H and S9L). In addition, the caspase-3 activation induced by *BCV* was substantially reduced in the absence of SLC9A9 (Figures 4I). During *BCV* infection in BMDMs, SLC9A9 dramatically localized to the fusion sites of LAMP1-associated vacuoles (Figures 4J). Thus, SLC9A9 was found to play important roles in *BCV*-triggered floatptosis.

### OMVs and barrel-like proteins of *BCV* play important roles in the formation of cytoplasmic vacuolization

Given the difference in membrane puff-ups and blebbing produced by *BC* and *BCV* (Figure 1C), we examined the OMVs derived from both *BC* and *BCV*. Notably, the production of intact OMVs in *BCV* was dramatically increased compared with that in *BC* (Figures 5A and 5B). Furthermore, the OMVs derived from *BCV* also triggered cytoplasmic vacuolization and caspase-3 activation (Figures 5C-5E). Next, we performed MS analysis of bacterial proteins and detected high levels of PorV (an outer membrane shuttle protein for the T9SS), lipocalin, β-barrel protein, and adenylosuccinate lyase (ADSL) in the supernatants of *BCV*-infected BMDMs but not in that of *BC*-infected cells (Figure S10A and Table S2). However, the expression of *PorV*, *lipocalin*, *β-barrel*, and *ADSL* was comparable between *BCV* and *BC* (Table S1), indicating that OMVs derived from *BCV* might contribute to the enrichment of these proteins in the supernatants of *BCV*-infected BMDMs. Thus, we cloned *PorV*, *lipocalin*, *β-barrel*, and *ADSL* from the *BCV* genome into the expression vector and purified the recombinant proteins (Figure S10B).

**Figure 5.**
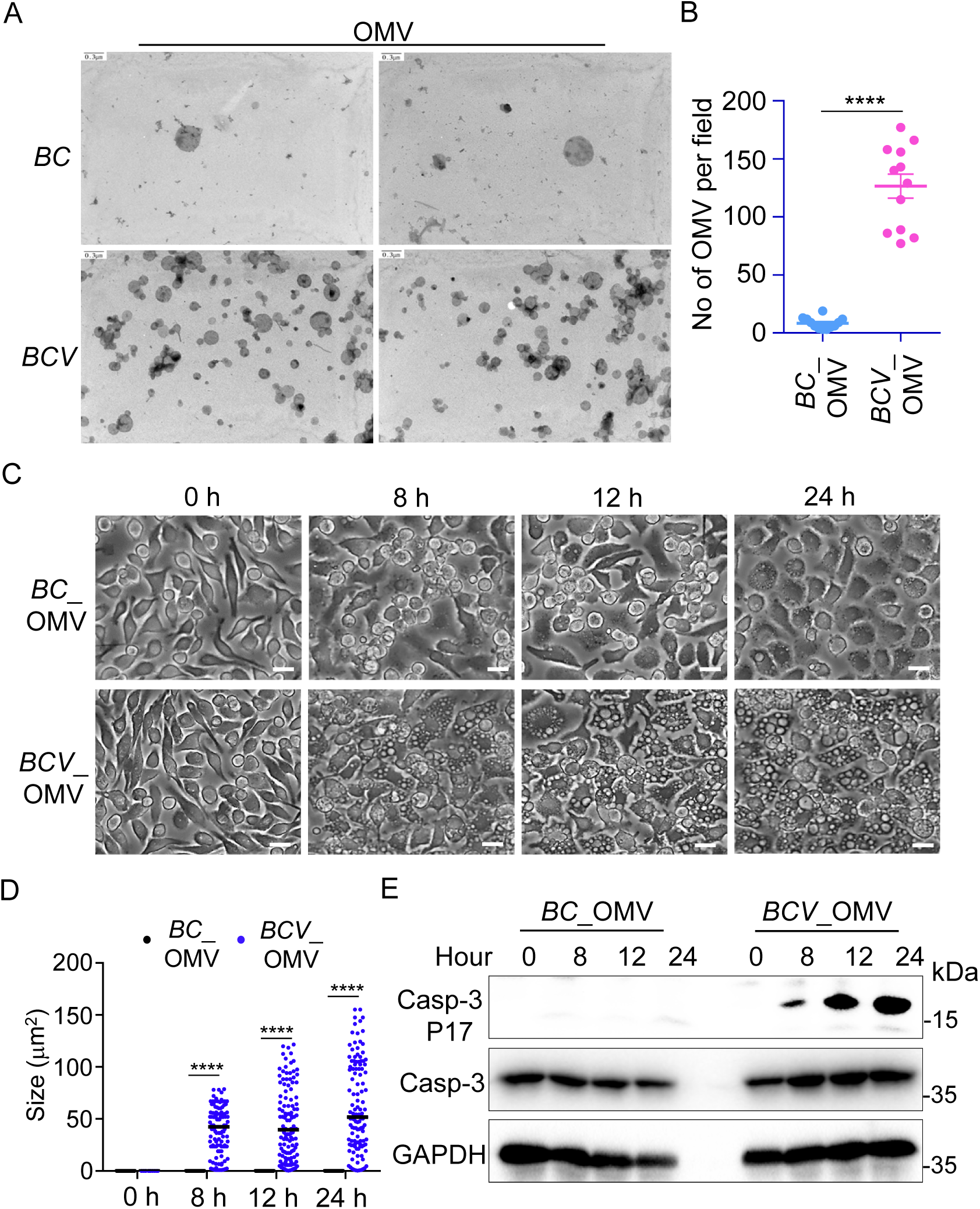
OMVs derived from *BCV* trigger the cytoplasmic vacuolization cell death. **A**, TEM analysis of purified OMVs from *BC* and *BCV*. *BC* and *BCV* were cultured on Columbia blood agar plates for 96 hours, followed by growth in BACTEC™ Lytic Anaerobic media for 16 hours. The supernatants of equal amounts of *BC* and *BCV* were used for OMV purification and comparative analysis. Scale bars, 0.3 μm. **B**, Quantification of the number of OMVs per field in (**A**) (n=12 random fields; 3 independent experiments). **C**, Microscopic analysis of WT BMDMs treated with OMVs derived from *BC* or *BCV* (100 μg) for the indicated times. Scale bars, 30 μm. **D**, Quantification of vacuole size in the BMDMs in (**c**). The largest vacuole per cell was analysed, and at least 120 cells were quantified for each group. **E**, Immunoblot analysis of caspase-3 and cleaved caspase-3 (P17) in WT BMDMs treated with OMVs derived from *BC* and *BCV* (100 μg) for the indicated times. Data are representative of 3 independent experiments with similar results (**A**-**E**). Data represent Mean ± SEM for (**B**, **D**), ****P < 0.0001, by 2-sided Student’s t test without multiple-comparisons correction.

The recombinant proteins of GST-fused PorV, GST-fused lipocalin, GST-fused β-barrel, GST-fused ADSL, or GST alone did not cause cytoplasmic vacuolization following their direct addition to the BMDMs (Figure S10C). However, transfection of GST-fused lipocalin or GST-fused β-barrel proteins into the cells resulted in profound cytoplasmic vacuolization between 1 and 12 hours after transfection (Figures 6A and S10D). The amount of cytoplasmic vacuolization caused by the transfection of the GST-fused PorV protein gradually decreased after 2 hours and substantially decreased up to 12 hours after treatment (Figures 6A and S10D). In contrast, transfection of GST-fused ADSL protein or GST protein alone did not cause cytoplasmic vacuolization in BMDMs (Figures 6A and S10D). The intracellular lipocalin, β-barrel, and PorV proteins were preferentially distributed around the fusion sites of LAMP1-associated vacuoles (Figure 6B). Moreover, transfection of the GST-fused lipocalin protein together with the GST-fused β-barrel protein or the combination of the GST-fused lipocalin, GST-fused β-barrel, and GST-fused PorV proteins induced the formation of large vacuoles at 12 hours and the formation of single vacuole-occupied cells at 20 hours after transfection (Figures 6C and S10E). Protein structure prediction revealed that lipocalin, β-barrel, and PorV all had multiple β-strands that formed barrel-like structures (Figure S10F). These results indicated that OMVs and the membrane proteins lipocalin, β-barrel, and PorV were important for *BCV*-induced floatptosis (Figure S11).

**Figure 6.**
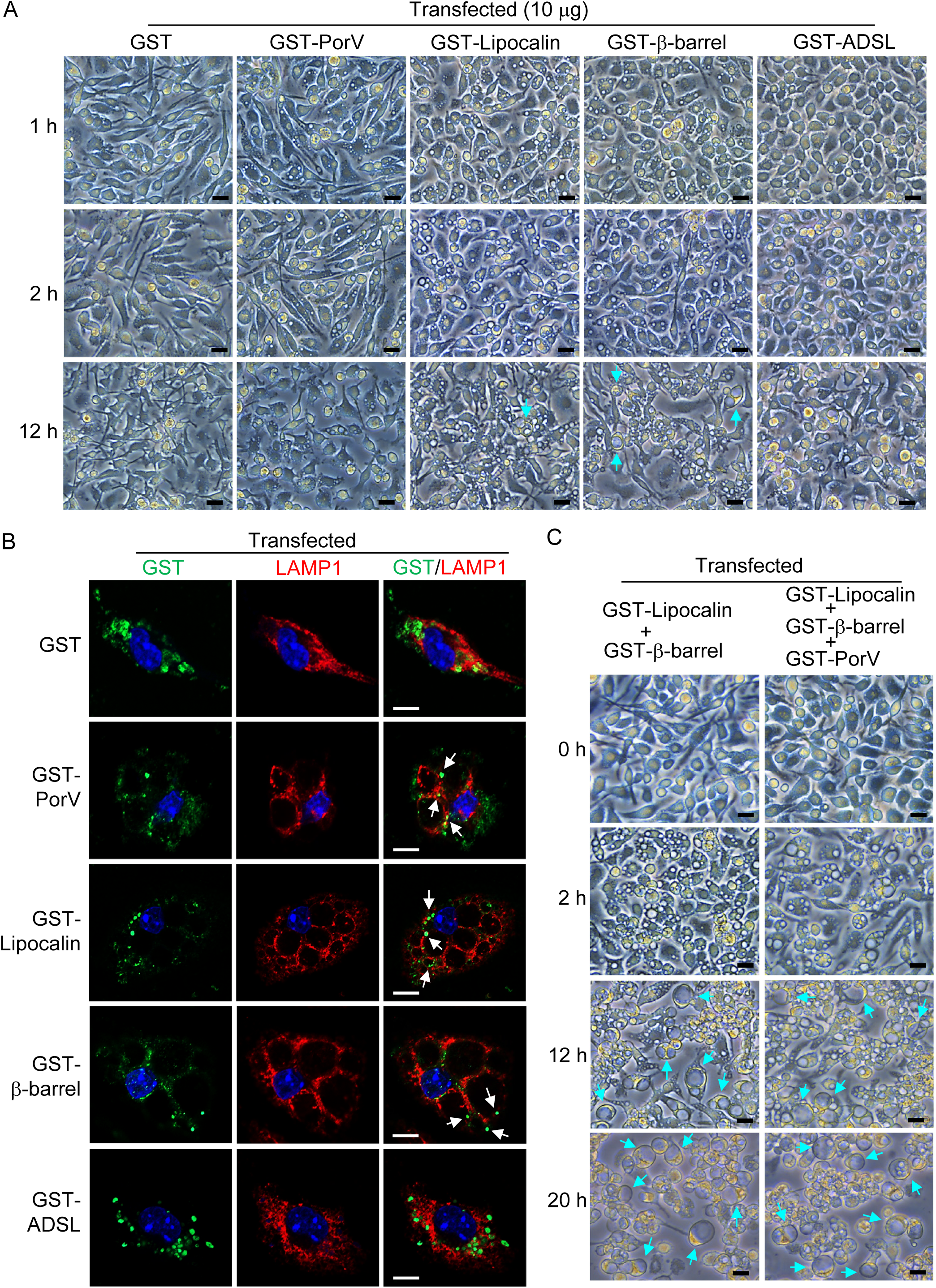
Transfection of the lipocalin, β-barrel, and PorV proteins of *BCV* induce the formation of cytoplasmic vacuolization. **A**, Microscopic analysis of WT BMDMs transfected with GST (10 μg), GST-PorV (10 μg), GST-Lipocalin (10 μg), GST-β-barrel (10 μg), and GST-ADSL (10 μg) proteins for 1, 2, and 12 hours as indicated. The arrows indicate large vacuole-containing cells. Scale bars, 30 μm. **B**, Confocal microscopy analysis of GST and LAMP1 in BMDMs transfected with GST (10 μg), GST-PorV (10 μg), GST-Lipocalin (10 μg), GST-β-barrel (10 μg), and GST-ADSL (10 μg) proteins for 2 hours. The arrows indicate the subcellular localization of the GST-fused proteins. Scale bars, 10 μm. **C**, Microscopic analysis of WT BMDMs transfected with the combination of GST-Lipocalin (10 μg), GST-β-barrel (10 μg), and GST-PorV (10 μg) proteins for the indicated times. The arrows indicate vacuole-occupied cells. Scale bars, 30 μm. Data are representative of 3 independent experiments (**A**-**C**).

### *BCV* increases in vivo pathogenicity

A bacterial killing assay revealed that the bacterial entry of *BC* into BMDMs was significantly lower than that of *BCV*, and the bacterial burden of *BCV* but not *BC* was dramatically greater at 22 hours post infection than at 2 hours post infection (Figure 7A). However, the bacterial growth of *BC* and *BCV* was comparable in culture media (Figure 7B), suggesting that *BCV* was more likely to undergo intracellular replication. To examine the pathogenicity of *BCV* and *BC* in vivo, we first infected alveolar macrophages (AMs) isolated from pulmonary lavage fluid ex vivo with *BC* or *BCV*. Consistent with the results obtained using BMDMs, *BCV*-infected AMs demonstrated remarkable cytoplasmic vacuolization (Figure 7C). Next, WT mice were intranasally infected with *BCV* or *BC* (4.0 × 10^8^ CFU per mouse), and the bacterial burden and host response were assessed. Both the body weight loss and bacterial burden in the lungs of the *BCV*-infected mice were significantly greater than those in the lungs of the *BC*-infected mice at 24 hours post infection (Figures 7D and 7E). Indeed, cytoplasmic vacuolization in AMs was observed in *BCV*-infected mice (Figure 7F). Furthermore, inflammatory responses and lung pathology, including vascular muscle hypertrophy, infiltration of immune cells, and inflammatory cytokine expression, were more severe in the *BCV*-infected than in the *BC*-infected mice (Figures 7G-7I).

**Figure 7.**
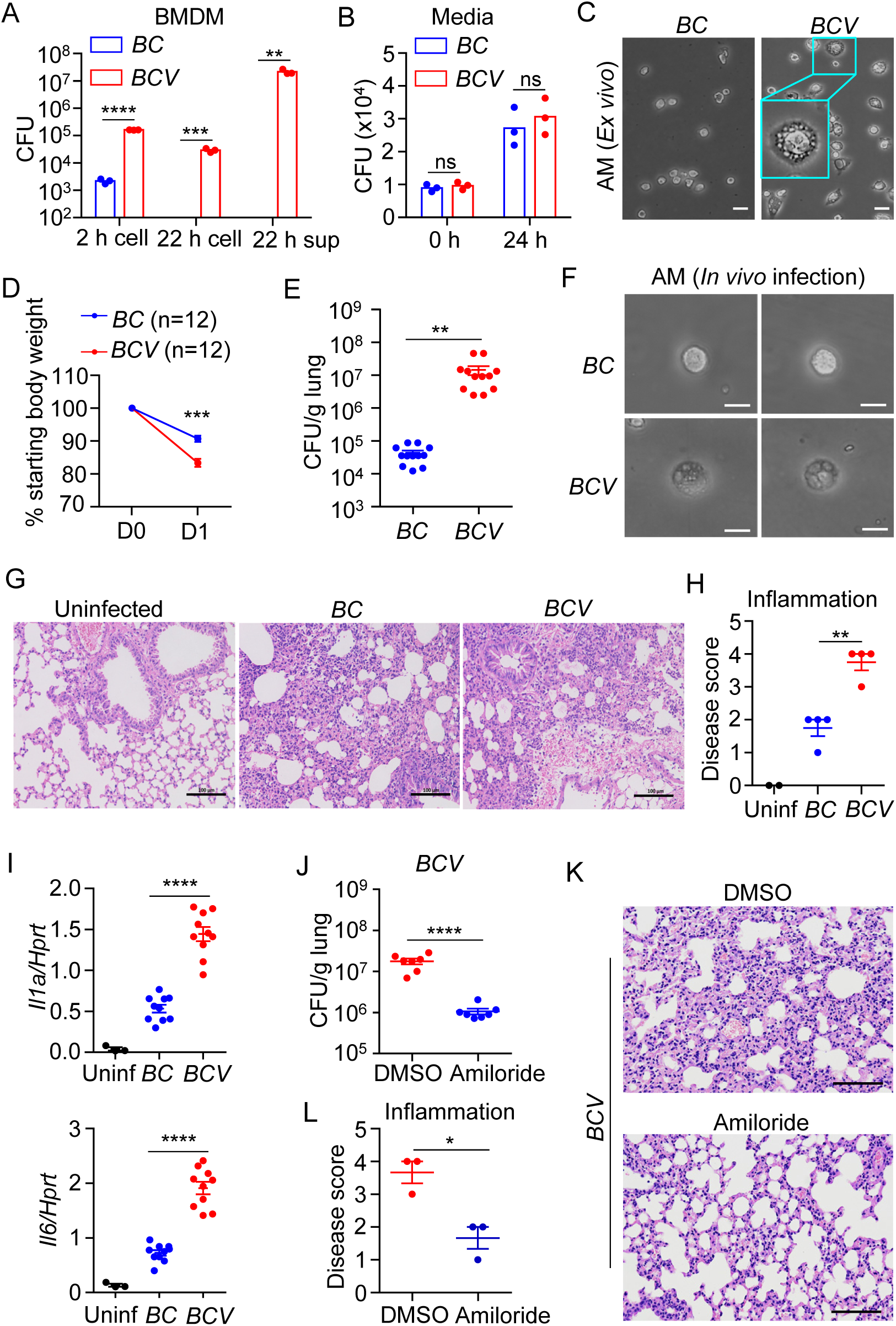
*BCV*-triggered cytoplasmic vacuolization cell death increases pathogenicity *in vivo*. **A**, Bacterial killing ability of *BC* and *BCV* in BMDMs. WT BMDMs were infected with *BC* (10 MOI) or *BCV* (10 MOI) for 2 hours. Infected BMDMs were washed, lysed and cultured on Columbia blood agar plates for 96 hours for enumeration of intracellular (2 h cell) bacteria. Washed BMDMs were further cultured in fresh media for 22 hours, and the numbers of intracellular (22 h cell) and extracellular (22 h Sup) bacteria were enumerated after they were cultured on Columbia blood agar plates for 96 hours (n=3 biologically independent samples). **B**, Growth analysis of *BC* and *BCV* in BMDM culture media. ‘0 h’ indicates the starting point of *BC* and *BCV*, and ‘24 h’ indicates the number of *BC* and *BCV* in the media after 24 hours of growth (n=3 biologically independent samples). **C**, Microscopic analysis of cytoplasmic vacuolization in Alveolar macrophages (AMs) infected with *BC* or *BCV* (400 MOI) for 12 hours. Scale bars, 20 μm. **D**,**E**, WT mice were intranasally infected with 4.0×10^8^ CFU *BC* (n=12) or *BCV* (n=12), and the body weight change (**D**) and bacterial burden in the lungs on Day 1 after infection were measured (**E**). **F**, Microscopic analysis of cytoplasmic vacuolization in isolated AMs from *BC*- and *BCV*-infected mice in (**E**). Scale bars, 20 μm. **G**, H&E staining of lung sections from uninfected and *BC*- and *BCV*-infected mice in (**E**). Scale bars, 100 μm. **H**, Disease scores on the basis of inflammation in lung sections in (**G**) from uninfected (Uninf, n=2), *BC*-infected (n=4), and *BCV*-infected (n=4) mice. **I**, Expression of genes encoding IL-1α and IL-6 was analysed in lung tissues from uninfected (Uninf, n=3) and *BC*-infected (n=10), and *BCV*-infected (n=10) mice in (**E**). **J**, WT mice were intranasally infected with *BCV* (4.0×10^8^ CFU), and the bacterial burden in the lungs was measured at 24 hours after infection. Amiloride indicates that the mice were intravenously injected with amiloride hydrochloride (10 mg/kg) twice, at 0 and 12 hours after *BCV* infection (n=7 mice for each group). **K**, H&E staining of lung sections from DMSO- or amiloride hydrochloride-treated mice infected with *BCV* in (**K**). Scale bars, 100 μm. **L**, Disease scores on the basis of inflammation in the lung sections in (**K**) (n=3 mice for each group). Data are from 3 independent experiments (**A**, **B**) or representative of 3 independent experiments with similar results (**C**-**K**). Data represent Mean ± SEM for (**A**, **B**, **D**, **E**, **H**, **I**, **J**, **L**), 2-sided Student’s t test without multiple-comparisons correction, *P < 0.05, **P < 0.01, ***P < 0.001, ****P < 0.0001.

Given that amiloride has the capacity to inhibit cytoplasmic vacuolization cell death induced by *BCV* in BMDMs, to determine whether cytoplasmic vacuolization contributed to the high pathogenicity of the *BCV* pathogen, we combined *BCV* administration with amiloride treatment in vivo, which might inhibit the cytoplasmic vacuolization induced by *BCV* infection. Interestingly, the bacterial burden of *BCV* in the lung was significantly reduced in the mice treated with amiloride (Figure 7J). In addition, H&E staining revealed that the lung pathology was reduced in the presence of amiloride (Figures 7K and 7L). Overall, these results indicated that *BCV* was a more virulent variant of the *Bergeyella cardium* bacterial strain and that *BCV*-triggered floatptosis enhanced its pathogenicity.

## Discussion

*Bergeyella cardium* is an emerging pathogen, and studies on *Bergeyella cardium* strains are mostly limited to clinical case reports^7,15^. In this study, we isolated *BCV* and *BC* from a patient with infectious endocarditis who had used antibiotics for a long time before the clinical examination^7^. Our work demonstrated that *BCV* exhibited several characteristics contributing to immune evasion, including acquired intracellular replication capacity, robust OMV release, increased serum killing resistance, downregulated inflammatory cytokine expression, induced cytoplasmic vacuolization cell death, and increased in vivo pathogenicity. Whether the long-term use of antibiotics in patients induced genetic changes or epigenetic switching of *BCV* to promote immune evasion capacity remains unclear.

Bacterial OMVs play a critical role in host–microbial interactions that influence pathogenesis through the delivery of virulence factors and the elicitation of inflammatory responses^16^. The large number of intact OMVs shed by *BCV* might result in delivery of the virulence factors lipocalin, β-barrel, and PorV into BMDMs to cause cytoplasmic vacuolization. Lipocalin family members are ancestral proteins that are found in all kingdoms of life, and secreted lipocalin polypeptides have been reported to induce apoptosis of leukocytes through an autocrine pathway^17^. β-barrel proteins contribute to the formation of β-barrel outer membrane proteins and are conserved across species in terms of folding and insertion into the outer membrane^18^. Transmembrane β-barrel proteins can fold spontaneously and assemble into lipid membranes to form stable pores^19^. The barrel-like structures of lipocalin, β-barrel, and PorV of *BCV* might contribute to their assembly into the membrane, leading to the initiation of cytoplasmic vacuolization and subsequent membrane fusion of vacuoles. The interplay between lipids and proteins plays a crucial role in dynamic membrane remodelling, and phosphatidylinositol bisphosphate (PIP_2_) controls the formation and spatiotemporal organization of protein complexes involved in vesicle budding, trafficking, and membrane curvature and fusion^20^. Interestingly, deficiency of the PIKfyve lipid kinase complex components, including the phosphoinositide (PI) kinase PIKfyve, the scaffolding protein Vac14, and the lipid phosphatase Fig4, inhibited the conversion of phosphatidylinositol-3-phosphate (PI3P) to phosphatidylinositol 3,5-bisphosphate PI(3,5)P_2_ and caused loss of PI(3,5)P_2_ and remarkable vacuolation in multiple cells in both mice and humans ^21–27^. Thus, the barrel-like proteins of *BCV* might regulate the homeostasis and distribution of PIP_2_ to modulate membrane dynamics and cytoplasmic vacuolization, which requires further investigation.

Genomic analysis revealed differences in genes between *BC* and *BCV*, including *PorV* and *BatD*. The Bat proteins have been shown to partially compensate for the oxidative stress response in bacteria and to provide defence against oxidative damage^28^. In *Francisella tularensis*, the BatD homologue mutant reduced the intracellular replication capacity in macrophages and virulence in a mouse model^29^. Thus, the addition of the BatD protein to *BCV* might partially contribute to stress adaptation, intracellular replication, and in vivo pathogenicity mechanisms. PorV is an important machinery component of T9SS and functions as a shuttle protein to deliver T9SS substrates to the attachment complex on the cell surface^30^. Structural prediction revealed that the deletion corresponding to residues 344–351 of *BCV* PorV led to structural changes in PorV. It is possible that the ‘*BCV* β15’ strand promoted β-barrel formation altogether, which might be associated with the robust production of OMVs in *BCV* through mediating the secretion and attachment of cargo proteins to the cell surface and OMVs^31,32^. We speculate that the increased virulence of *BCV* could be partially due to the robust production of OMVs, which penetrate host cells, causing cytoplasmic vacuolization and cell death. A limitation of this study is the lack of genetic knockout and complementation experiments in *Bergeyella cardium* strains to confirm whether these genomic mutations in *BCV* contributed to robust OMV biogenesis and the secretion of virulence factors. Challenges remain in establishing genetic manipulation techniques for novel organisms due to the lack of easy-to-handle genetic manipulation tools^33^, which should be a focus for future development.

Cytoplasmic vacuolization is an evolutionarily conserved event that plays dual roles in cell death and survival depending on transient or irreversible vacuolization properties^6^. The equilibration of osmotic pressure within organelles by water diffusion, rather than programmed gene expression and protein regulation, is involved in the process of transient vacuolization formation induced by natural or synthetic chemical compounds^34,35^. In this study, we found that amiloride (an inhibitor of Na^+^/H^+^ exchange) was able to inhibit the floatptosis induced by *BCV* infection, indicating that a transmembrane pH gradient was required for this vacuolization process. Excess autophagy-induced autosis cell death was previously reported to depend on vacuolar Na^+^, K^+^-ATPase activity^36^, suggesting the important role of the pH gradient in endosomal–lysosomal fusion and cytoplasmic vacuolization. More importantly, we identified the solute carrier family 9 member SLC9A9 as a gatekeeper for *BCV*-triggered floatptosis.

Solute carriers are a large group of membrane transport proteins with more than 300 members in humans, and the SLC9 family (also called sodium/hydrogen exchanger 9, NHE9) is divided into three subgroups on the basis of membrane localization and cell type specificity^37^. SLC9A9 colocalizes with intracellular late endosomes and contributes to the transport of sodium and hydrogen ions across the membrane to maintain the pH balance of intracellular organelles^37–39^. Overexpression of SLC9A9 has been reported to lead to endosomal alkalization, SLC9A9 knockdown has been shown to induce endosomal acidification, and the organellar pH is essential for enzyme activity, membrane fusion, and cell volume; however, the detailed mechanism is not clear^37^. The expression level of *Slc9a9* induced by *BCV* was much greater than that triggered by *BC*, suggesting that *BCV* maintains high expression of *Slc9a9* to prevent acidification of vacuoles and promote vacuole fusion for intracellular survival. However, the interplay between SLC9A9 and the intracellular PIP_2_ composition and localization that regulate membrane fusion remains elusive. In addition to SLC9A9, other ATPases and ion channels might also be involved in the process of *BCV*-induced cytoplasmic vacuolization. Thus, SLC9A9 represents one of the major targets modulated by *BCV* to hijack for immune evasion.

Solute carrier transporters are important metabolic regulators of immune cells and play critical roles in cancer immunotherapy^40,41^. Dysfunction of SLC9 family members is associated with multiple diseases, such as cancer, neurological disorders, gastrointestinal tract disorders, and kidney disease^42–44^. The vacuolization of hematopoietic precursors has been recently recognized as a feature of inflammatory syndrome, but the molecular mechanisms involved in this pathological process are not known^45^. Thus, the finding of SLC9A9-mediated cytoplasmic vacuolization cell death upon *BCV* infection provides novel insights into the mechanisms of both SLC9A9- and vacuolization-associated diseases.

## EXPERIMENTAL PROCEDURES

### Mice

*Ifnar^-/-^*, *Aim2^-/-^Nlrp3^-/-^*, and *Asc^-/-^* mice were previously described^46^. *Slc9a9^-/-^* mice were generated by GemPharmatech Co., Ltd (Suzhou, Jiangsu). Exon 2 of the *Slc9a9* gene was knocked out by CRISPR-Cas9 system. The strategy for construction of the targeting vector is illustrated in Figure S9i. The knockout of *Slc9a9* gene was validated by genotyping, qRT-PCR and western blot (Figs. S9J, S9K and 4I). WT and knockout mice were kept under specific pathogen-free conditions in the Animal Resource Center at Shandong University, Jinan, Shandong Province, China. All animal experiments were conducted in accordance with guidelines approved by the Ethics Committee of Scientific Research of Shandong University.

### Bacteria and OMVs Preparation

*BC* and *BCV* were cultured on Columbia blood agar plates (Autobio,0004763) for 96 hours, followed by growing in BACTEC™ Lytic Anaerobic media (BD, 442021) with shaking (220 rpm) for 16 hours at 37 °C. The bacteria were precipitated by centrifugation at 3,000 rpm for 30 minutes at 4 °C, and resuspended in DMEM/F-12 media for infection experiments. For OMVs preparation, the supernatants were passed through a 0.45 µm filter to remove debris, and the filtered supernatants were precipitated by ultracentrifugation at 40,000 g for 2 hours at 4°C, and resuspended in BACTEC™ Lytic Anaerobic media and BMDM-growing media for TEM analysis and treatment experiments, respectively.

### Preparation of BMDMs, treatment, bacterial infection, and siRNA transfection

To generate BMDMs, bone marrow cells were cultured in L929 cell-conditioned DMEM/F-12 supplemented with 10% FBS, 1% nonessential amino acids, and 1% penicillin-streptomycin for 5 days. The siRNAs target to *Slc9a9*, *Slc9a1*, *Slc9b2*, *Slc12a2*, *Slc13a3*, *Slc35g1*, *Atp1a1*, *Rab7*, *Gpr65*, *Cd5l*, *Olfr56*, *Ptpn22*, and *Tlr9* were ordered from RiboBio Company. siRNAs were electroporated into BMDMs using the Neon^TM^ Transfection System following the manufacturer’s instructions. The siRNA sequences are listed in Supplementary Table 4. Inhibitors z-VAD (Calbiochem, 627610), necrostatin-1 (Nec-1; Calbiochem, 480065), necrosulfonamide (NSA; MCE, HY-100573), ferrostatin-1 (Fer-1; MCE, HY-100579), pyrrolidinedithiocarbamate ammonium (PDTC; TargetMoI, T3147), 3-Methyladenine (3-MA; APExBIO Technology, A8353), wortmannin (CST, 9951S), rapamycin (MCE, HY-10219), and amiloride hydrochloride (Alomone labs, A-140) were used to treat BMDMs for 2 hours with indicated concentration ahead of bacterial infection.

To induce floatptosis, WT BMDMs in 12-well plate (1 M per well) were infected with different MOIs of *BCV* (100-400 MOIs) up to 40 hours, *BCV*-infected BMDMs were examined by microscopy and cytoplasmic vacuolization was evaluated according the number and size of vacuoles by ImageJ software. We found that *BCV* at 400

MOI can induce more cytoplasmic vacuolization and large vacuoles at 12 hours and 24 hours post infection compared with lower MOIs. Thus, 400 MOI was used for in vitro infection experiments. For comparing experiments, WT, inhibitor-treated, and siRNA-transfected BMDMs were infected with *BC* and *BCV* (400 MOI) as indicated times. The treated and control cells were analyzed for cytoplasmic vacuolization under a microscope and lysed for RNA and protein analysis.

### Bacterial killing assay

BMDMs were infected with *BC* and *BCV* with a MOI of 10 for 2 hours and washed; cells were washed twice and cultured in fresh media (DMEM/F-12 and 10% FBS) without antibiotic. After 20 hours, the supernatant (extracellular bacteria) and BMDMs lysed in PBS (intracellular bacteria) were serially diluted, and plated onto Columbia Blood agar plates, and incubated 96 hours for CFU enumeration.

### Bacterial infection of mice

The bacterial strains *BC* and *BCV* were cultured on Columbia blood agar plates for 96 hours, followed by growing in BACTEC™ Lytic Anaerobic media (BD, 442021) with shaking (220 rpm) for 16 hours at 37 °C. Eight-to ten-week-old and same gender wild type mice were infected intranasally with *BC* and *BCV* (4.0×10^8^ CFUs per mouse). Mice were weighed and monitored at day 0 and day 1. Mice were euthanized at day 1 after infection and lungs were harvested to determine the bacterial burden and cytokine expression.

### Immunoblot analysis and antibodies

Samples were separated by 12% SDS-PAGE, followed by electrophoretic transfer to polyvinylidene fluoride membranes, and membranes were blocked and then incubated with primary antibodies. The following primary antibodies were used: anti-caspase-3 (CST, 9662); anti-cleaved-caspase-3 (CST, 9661); anti-SLC9A9 (Proteintech, 66577-1-Ig; Invitrogen, PA5-101895); anti-caspase-1 (AdipoGen, AG-20B-0042); anti-MLKL (Abcepta, AP14272B); anti-p-MLKL (CST, 37333); anti-RIP3 (CST, 95702S); anti-p-RIP3 (CST, 91702S); anti-p-PI3K (CST, 4228); anti-p-S6 (CST, 4856); anti-Cas9 (abcam, ab191468) and anti-GAPDH (CST, 5174). HRP-labeled anti-rabbit (CST, 7074), anti-mouse (CST, 7076), or anti-goat (Santa Cruz, sc-2006) was used as the secondary antibody.

### Immunofluorescence staining and microscopy

For LAMP1, RAB7, SLC9A9, GST, EEA1, and RAB5 immunostaining, treated and untreated BMDMs were fixed in 4% paraformaldehyde for 15 minutes at room temperature. Cells were washed with PBS and blocked in 1×ELISA buffer with 0.1% saponin for 1 hour. Cells were stained with anti-LAMP1 (Invitrogen, 14-1071-85); anti-RAB7 (CST, 9367); anti-SLC9A9 (Proteintech, 66577-1-Ig), anti-GST (Proteintech, 10000-0-AP), anti-EEA1 (CST,3288) or anti-RAB5 (CST, 3574)— all at 1:300 to 1:500 dilution, overnight at 4 °C. Cells were washed, stained with a fluorescence conjugated secondary antibody (Invitrogen, A-11008, Alexa Fluor™ 488, Goat anti-Rabbit; Invitrogen, A-21422, Alexa Fluor™ 555, Goat anti-Mouse; Invitrogen, A-11077, Alexa Fluor™568, Goat anti-Rat) at 1:300 dilution for 40 minutes at 37 °C, and mounted using a mounting medium (Vector Laboratories, H-1200). Lucifer Yellow was ordered from Aladdin (L131282). Cells were observed on the ZEISS-LSM880 and ANDOR High speed confocal microscope, image acquisition and data analysis were performed using ZEN black_2-3SP1, ZEN blue 2.6, and Imaris software.

### Live-cell imaging for cell death

BMDMs (0.5×10^6^ cells/well) were seeded in 24-well plates. BMDMs were infected with *BC* and *BCV*, and stained with propidium iodide (PI; Life Technologies, P3566) and Hoechst (Beyotime Biotechnology, C1029) according to the manufacturer’s instruction. The plate was scanned and images were acquired in real-time every 2 hours from 0 to 24 h post-treatment by Opera Phenix High Content Screening System by PerkinElmer, Inc. Hoechst staining indicates the total number of cells, PI-positive dead cells are marked with a red mask for visualization. The image analysis, masking, and quantification of dead cells were done using the Harmony software package supplied with the system.

### RNA sequencing and data analysis

Total RNA was extracted from *BC*- and *BCV*-infected and uninfected WT BMDMs cultured in L929 cell-conditioned DMEM/F-12 media and subjected to commercial RNA-sequencing (RNA-seq) analysis (Novogene). The RPKM values (reads per kilobase of transcript per million reads mapped) for each transcript were calculated. Differential expression analysis between comparative groups was conducted using DESeq2 software (version 1.42.1). Differentially Expressed Genes (DEGs) were selected based on the criteria of |log2(Fold change)| ≥ 1 and a p-value less than 0.05. KEGG pathway enrichment analysis of the DEGs were performed using clusterProfiler software (version 4.10.1).

*BC* and *BCV* were cultured on Columbia blood agar plates (Autobio,0004763) for 96 hours, and the bacterial RNA was extracted for RNA expression analysis (LC-BIO Biotech). Differential expression analysis was performed with R package edgeR (version 4.2.0). DEGs were selected based on a logarithmic fold change greater than 2 and a false discovery rate (FDR) below 0.05. To understand the functions of the DEGs, KEGG pathway analyses were conducted using KOBAS. DEGs were considered significantly enriched if their Bonferroni-corrected p-value was less than 0.05. The heat map was plotted using the R package pheatmap version 1.0.12.

### Library preparation, genome sequencing and assembly

The whole genome of *BC* and *BCV* were sequenced using PacBio Sequel platform and Illumina NovaSeq PE150 at the Beijing Novogene Bioinformatics Technology Co., Ltd. Briefly, the DNA sample was fragmented by sonication to a size of 350 bp, then DNA fragments were end-polished, A-tailed, and ligated with the full-length adaptor for Illumina sequencing with further PCR amplification. At last, PCR products were purified (AMPure XP system) and libraries were analyzed for size distribution by Agilent2100 Bioanalyzer and quantified using real-time PCR. To improve analysis accuracy, the low-quality reads were filtered (less than 500 bp) to obtain Clean data. The long reads were selected by SMRT portal (more than 6,000bp) as the seed sequence, and the other shorter reads were aligned to the seed sequence by Blasr. The results of preliminary assembly were corrected by the SMRT Link software with Illumina data. Cyclization was confirmed according to the overlap between the head and the tail, and the initiation site was corrected by blast with the DNAa database. The genome component prediction and gene annotation were done by National Center for Biotechnology Information (NCBI). The dataset of genome sequence has been deposited in the GenBank under accession no. CP029149 and CP114055. The genomic alignment between two *BC* and *BCV* was conducted using SnapGene, and scalable circular genome maps were generated using the Java package CGView to display the alignment of the *BCV* genome with the *BC* genome.

### MS analysis

The supernatants from *BCV*- and *BC*-infected BMDMs were performed with Mass spectrometry (MS) analysis by Applied Protein Technology (Shanghai). Briefly, protein digestion was performed by trypsin. The digest peptides of each sample were desalted on C18 Cartridges, concentrated by vacuum centrifugation and reconstituted in 40 µl of 0.1% (v/v) formic acid. MS analysis was performed on a timsTOF Pro mass spectrometer (Bruker) that was coupled to Nanoelute (Bruker Daltonics) for 45 minutes. The peptides were loaded on a C18-reversed phase analytical column in buffer A (0.1% formic acid) and separated with a linear gradient of buffer B (99.9% acetonitrile and 0.1% formic acid) at a flow rate of 300 nl/min. The mass spectrometer was operated in positive ion mode. The MS raw data for each sample were combined and searched using the MaxQuant software for identification and quantitation analysis. Proteins that were identified in the supernatants from *BCV*- and *BC*-infected BMDMs are listed in Table S2.

### Plasmid construction

The full-length of *Slc9a9* was amplified from a mouse cDNA library and subcloned into the pCDH vector. *Lipocalin*, *β-barrel*, *ADSL*, and *PorV* genes were amplified from the DNA of *BCV* and subcloned into the pGEX-6P-2 vector. One gRNA was designed to target exon 4 of *Slc9a9* gene and the corresponding double stranded oligonucleotide was subcloned into the lentiCRISPR vector following the CRISPR/Cas9 design instruction. All plasmids were confirmed by DNA sequencing. The primer sequences for vector construction are listed in Table S4. Lipofectamine 3000 reagents were used for transient transfection of plasmids into HEK293T cells.

### Expression, purification, and transfection of recombinant proteins

The control PGEX-6P-2 vector and GST-fused protein constructs PGEX-6P-2-PorV, PGEX-6P-2-Lipocalin, PGEX-6P-2-β-barrel, and PGEX-6P-2-ADSL were transformed into the Escherichia coli strain Rosetta (DE3) pLysS for expression. The expressed proteins were loaded onto Glutathione Agarose (MCE) and Mono S^TM^ 5/50 GL (Cytiva, USA) pre-equilibrated with the lysis buffer for purification. Endotoxin was removed using Pierce High-Capacity Endotoxin Removal Resin (Thermo, 88274) according to the manufacturer’s protocol. GST and GST-fused proteins were transfected into the cells using Xfect™ Protein Transfection Reagent (TaKaRa, 631324) following the manufacturer’s instruction.

### Lentivirus production and infection

The viral particles were prepared by transfecting HEK293T cells with *Slc9a9* overexpression and knockout plasmids in combination with packaging vectors pMDLg/pRRE, pRSV-Rev, and pCMV-VSV-G for overexpression; psPAX2 and pCMV-VSV-G for knockout. Ten hours later, the Opti-MEM media were replaced with fresh complete DMEM. Viral supernatant was harvested and passed through 0.45 μm syringe filter 48 and 72 hours after transfection. To establish stably transfected cells, Bone marrow cells, and iBMDMs were infected 2 times with filtered lentiviral supernatant in the presence of polybrene (8 μg/ml). The transduced BMDMs and iBMDMs were cultured in fresh BMDM-growing media for generation or further treatments and analysis.

### Transmission Electron Microscopy (TEM) and Scanning Electron Microscopy (SEM)

*BC*- and *BCV*-infected BMDMs were fixed in 2% paraformaldehyde and 2.5% glutaraldehyde in 0.1 M cacodylate buffer (pH 7.4) for 1 hour at 37 °C. Cells were embedded and sectioned for TEM by Jindi Medical Technology (Jinan, Shandong). *BC* and *BCV* were cultured on Columbia blood agar plates for 96 hours, followed by growing in BACTEC™ Lytic Anaerobic media (BD, 442021) with shaking (220 rpm) for 16 hours at 37 °C. The equal number of bacteria were precipitated and fixed in 2.5% glutaraldehyde and phosphate buffer, and sectioned samples were analyzed by TEM and SEM at Servicebio (Wuhan, Hubei).

### Preparation of tissue sample for HE staining

The superior lobes of the right lungs were fixed in 10% formalin, and 5-μm sections were stained with hematoxylin and eosin (H&E) and examined with a microscope. The severity of lung disease was scored on the basis of the presence of inflammation by a pathologist blinded to the experimental groups according the grade standard (0 = absent; 1 = rare, minimal; 2 = scattered mild; 3 = multifocal, moderate; 4 = extensive, marked; 5 = severe).

### Real-time qRT-PCR

Total RNA was isolated from cells and tissues using TRIzol reagent (Invitrogen, Thermo Fisher Scientific). cDNA was reverse transcribed using M-MLV reverse transcriptase (Promega). Real-time qRT-PCR was performed on the Roche LightCycler 96 Real-Time Detection System. The primer sequences are listed in Table S4.

### Lactate dehydrogenase (LDH) release assay

Cell culture supernatants were collected at the indicated times, and lactate dehydrogenase activity was measured by using the Promega cytotoxicity kit (G1781) according to the manufacturer’s protocols.

### Flow cytometry analysis

For flow cytometric analysis of cell death, *BC*- and *BCV*-infected WT BMDMs were stained with PI (Life Technologies, P3566) and Annexin V (BioLegend, 640918) for 30 minutes according to the manufacturer’s protocols, and cells were analyzed on a BD LSR Fortessa Cell Analyzer (BD Biosciences).

### Protein structure prediction and analysis

Protein structure of Lipocalin, β-barrel, PorV, and ADSL were predicted by the online server trRosetta (https://yanglab.qd.sdu.edu.cn/trRosetta/)^12,47,48^, and visualized by UCSF ChimeraX^49^. Briefly, the protein sequence was used as input sequence for structure prediction, performed with default parameter settings. trRosetta provided 5 models as results for each prediction. The highest ranked model of each prediction, model 1, was used for further analysis. The confidence of the overall structure prediction is reflected by the TM-score. A TM-score (0-1) above 0.5 usually indicates a model with correct topology. The confidence of the prediction at each residue is indicated by per-residue LDDT scores, ranging from 0-100, locating at the B-factor column of the PDB file of the structure. For structure analysis and comparison of *BC* and *BCV* PorV, structural figures were made using Open-Source PyMOL (Schrödinger). *BCV* and *BC* PorV models are colored by ‘b-factor’ ‘spectrum’, reflecting the per-residue LDDT scores. The structure model of *BC* PorV was superposed onto *BCV* PorV using ‘super’ command in PyMOL, resulting in an RMSD of 0.489 over 1961 atoms.

### Statistics

Data are presented as the mean ± standard error of the mean (SEM). Statistical analyses were performed using two-way ANOVA, 2-tailed Student’s *t* and log-rank tests. *P*-values of 0.05 or less were considered significant.

### Study approval

The present study was approved by the Ethics Committee of Scientific Research of Shandong University (ECSBMSSDU2021-2-171 and ECSBMSSDU2021-1-086), Jinan, Shandong Province, China.

## Author contributions

XQ, RM, HP, YZ, and TX designed the study. RM, HP, LY, ZL, XY, YL, YC, YY, WW, CG, JP, TX, YZ, and XQ performed experiments and analyzed the data. XQ, TX, and YZ wrote the manuscript. The order of co–first authorship was determined by the authors’ contributions to data acquisition and drafting of the manuscript.

## Acknowledgments

We Thank Z. Li, C. Huang, Q. Feng (Shandong University) and X. He (The Forsyth Institute, Cambridge, MA) for providing technical assistance and discussion. This work was supported by the National Natural Science Foundation of China (82125021, 82072255, 82272410, 82321002 and 81972005), Cutting Edge Development Fund of Advanced Medical Research Institute (GYY2023QY01), Shandong Province (2022GJJLJRC02-005) and Taishan Scholar Program of Shandong Province of China (tstp20221156).

---

**Figure S1.**
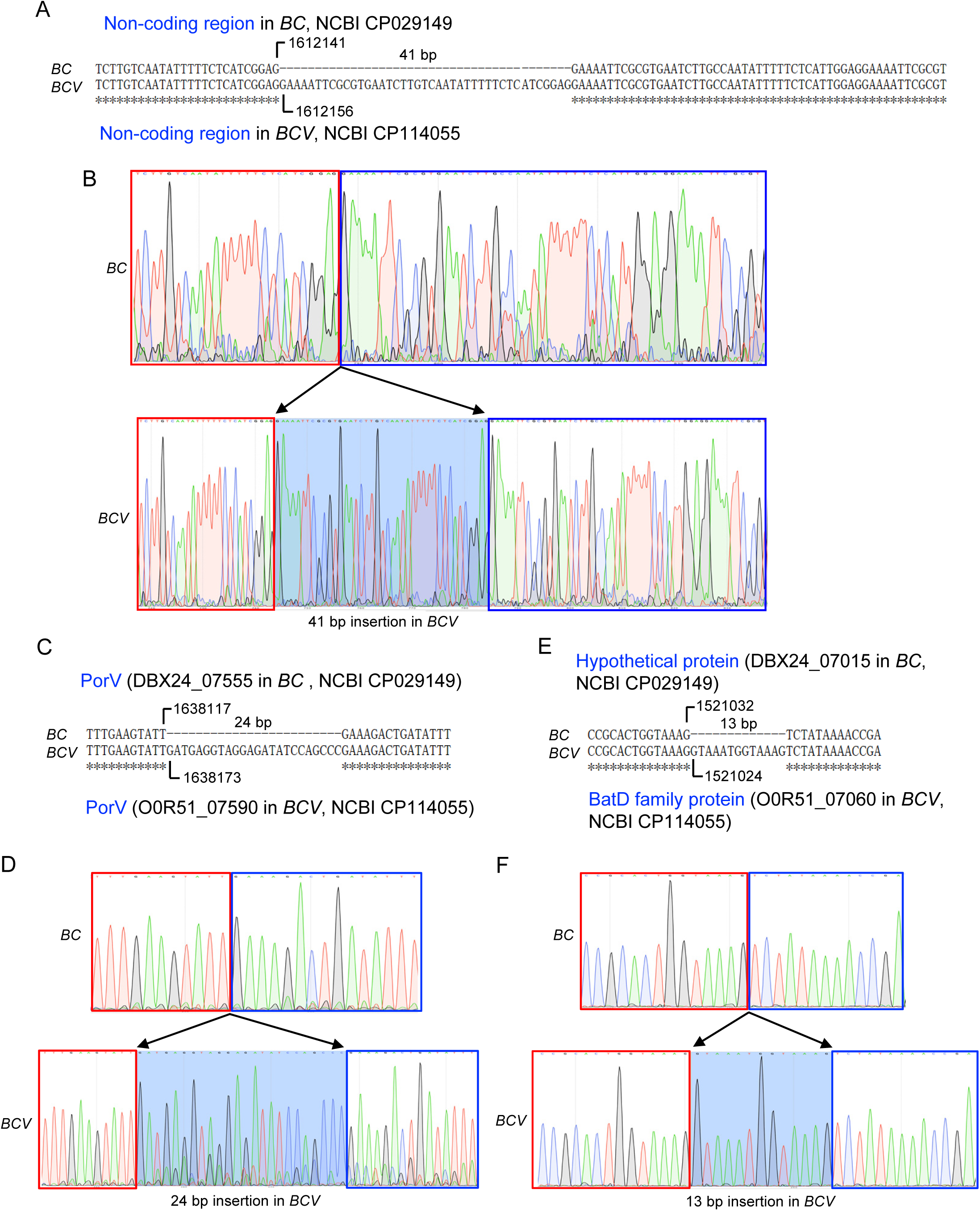
Validation of the genomic differences between *BC* and *BCV*. **A,** Sequence Alignment by CLUSTALW of non-coding region from the genomes of *BC* (CP029149) and *BCV* (CP114055). The number indicates the sequence position within genome. **B,** Validation of the difference in (**A**) by sanger sequencing. **C,** Sequence Alignment by CLUSTALW of *PorV* gene from the genomes of *BC* (CP029149) and *BCV* (CP114055). The number indicates the sequence position within genome. **D,** Validation of the difference in (**C**) by sanger sequencing. **E,** Sequence Alignment by CLUSTALW of hypothetical gene from the genomes of *BC* (CP029149) and *BCV* (CP114055). The number indicates the sequence position within genome. **F,** Validation of the difference in (**E**) by sanger sequencing.

**Figure S2.**
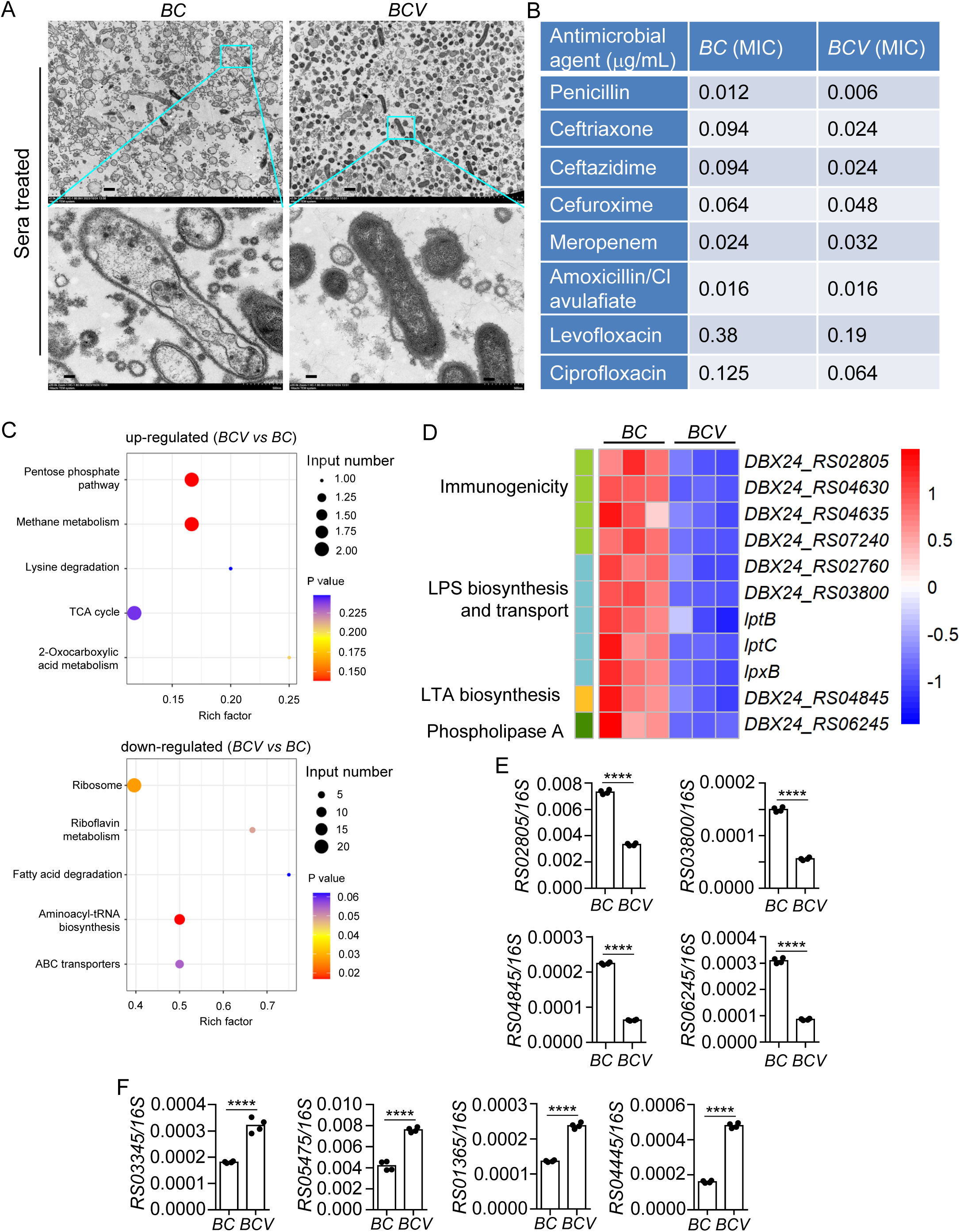
Antibiotic resistance and gene expression analysis of *BC* and *BCV*. **A**, TEM analysis of *BC* and *BCV* bacteria treated with human sera for 2 hours. Sera treated *BC* and *BCV* were precipitated, fixed, and sectioned for TEM analysis. Scale bars, 10 μm for upper and 1 μm for lower. **B**, Minimum inhibitory concentration (MIC) analysis of *BC* and *BCV* in the presence of different antimicrobial agents. **C**, KEGG enrichment analysis of differentially expressed genes between *BCV* and *BC*. The top 5 enriched metabolic pathways of upregulated and downregulated genes (*BCV* vs *BC*) are displayed. The size of each dot represents the number of DEGs, and the color gradient from blue to red indicates the significance level (p-value) of the enrichment analysis. **D**, Heatmap analysis of genes in (**C**) that are highly expressed in *BC* but not in *BCV*. **E**, Quantitative RT-PCR analysis of *RS02805*, *RS03800*, *RS04845*, and *RS06245* expression in *BC* and *BCV* strains (n=4 technical replicates; 3 independent experiments). *16S* rRNA was used as internal control to normalize the bacterial gene expression. **F**, Quantitative RT-PCR analysis of *RS03345*, *RS05475*, *RS01365*, and *RS04445* expression in *BC* and *BCV* strains (n=4 technical replicates; 3 independent experiments). *16S* rRNA was used as internal control to normalize the bacterial gene expression. Data are **from** 3 independent experiments (**C**, **D**) or representative of 3 independent experiments with similar results (**A**, **B**, **E**, **F**). Data represent Mean ± SEM for (**E**, **F**), ****P < 0.0001, by 2-sided Student’s t test without multiple-comparisons correction.

**Figure S3.**
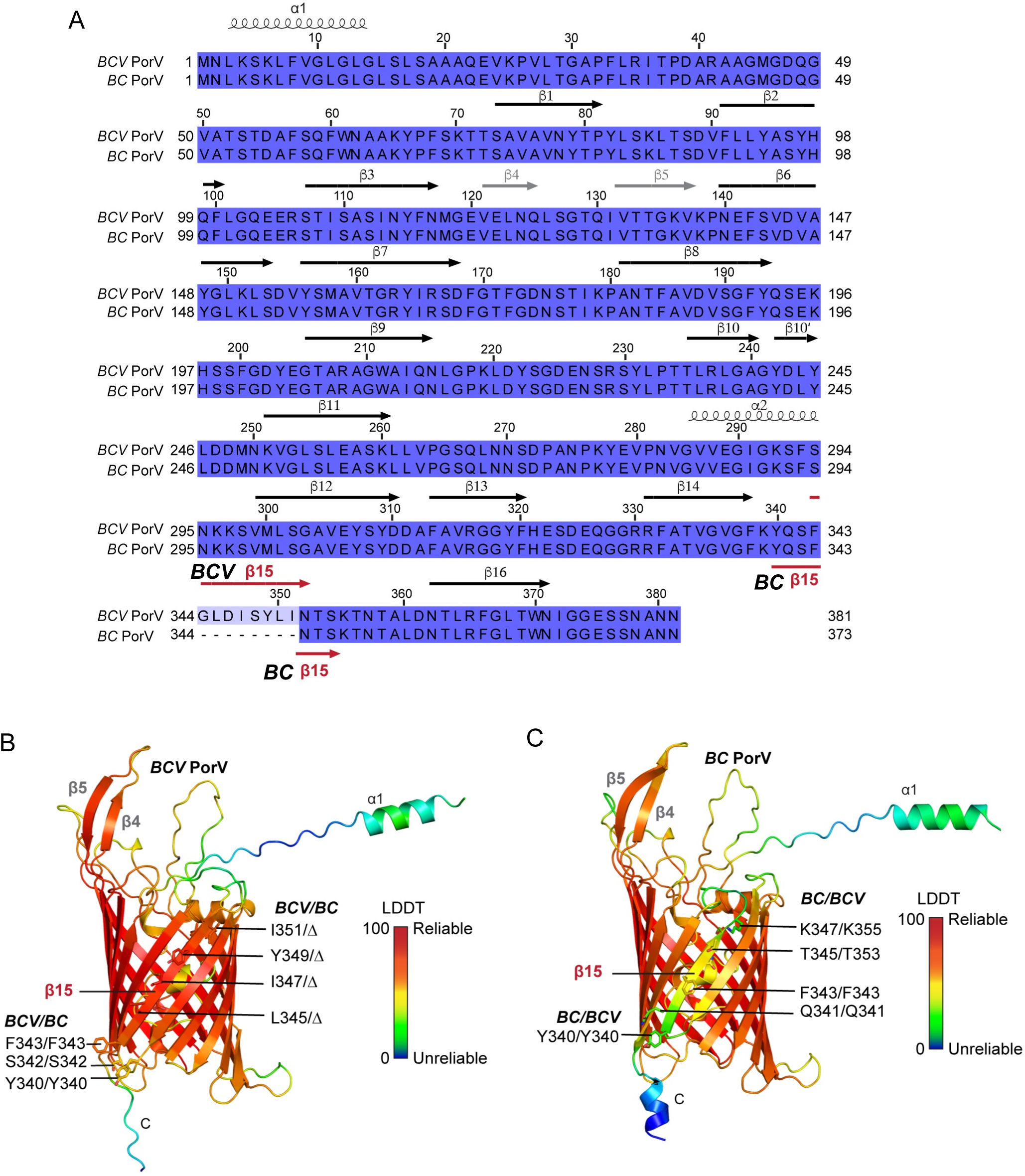
Sequence alignment and structure analysis of PorV. **A**, Sequence alignment of *BCV* and *BC* PorV. The secondary structure of *BCV* PorV is indicated on top of the alignment, based on the predicted structure of *BCV* PorV. Helices indicate α-helices in the structure. The arrows indicate β-strands. The *BC* β15 PorV is indicated below the sequence alignment. **B**, Structural model of the *BCV* PorV predicted by trRosseta. The structure is colored with spectrum from blue to red, according to LDDT values (0-100), which reflects the reliability of the prediction at each residue. The overall estimated TM-score is 0.915. The ‘*BCV* β15’ is labeled as β15, and the sidechains for ‘*BCV* β15’ and nearby residues are shown as sticks. The residue numbers of *BCV* and *BC* PorV are both indicated for comparison. β means the residue is deleted. **C**, Structural model of the *BC* PorV predicted by trRosseta. The model was superposed onto *BCV* PorV and shown in the same orientation as *BCV* PorV. The color scheme is the same as in (**B**). The overall estimated TM-score for *BC* PorV is 0.899.

**Figure S4.**
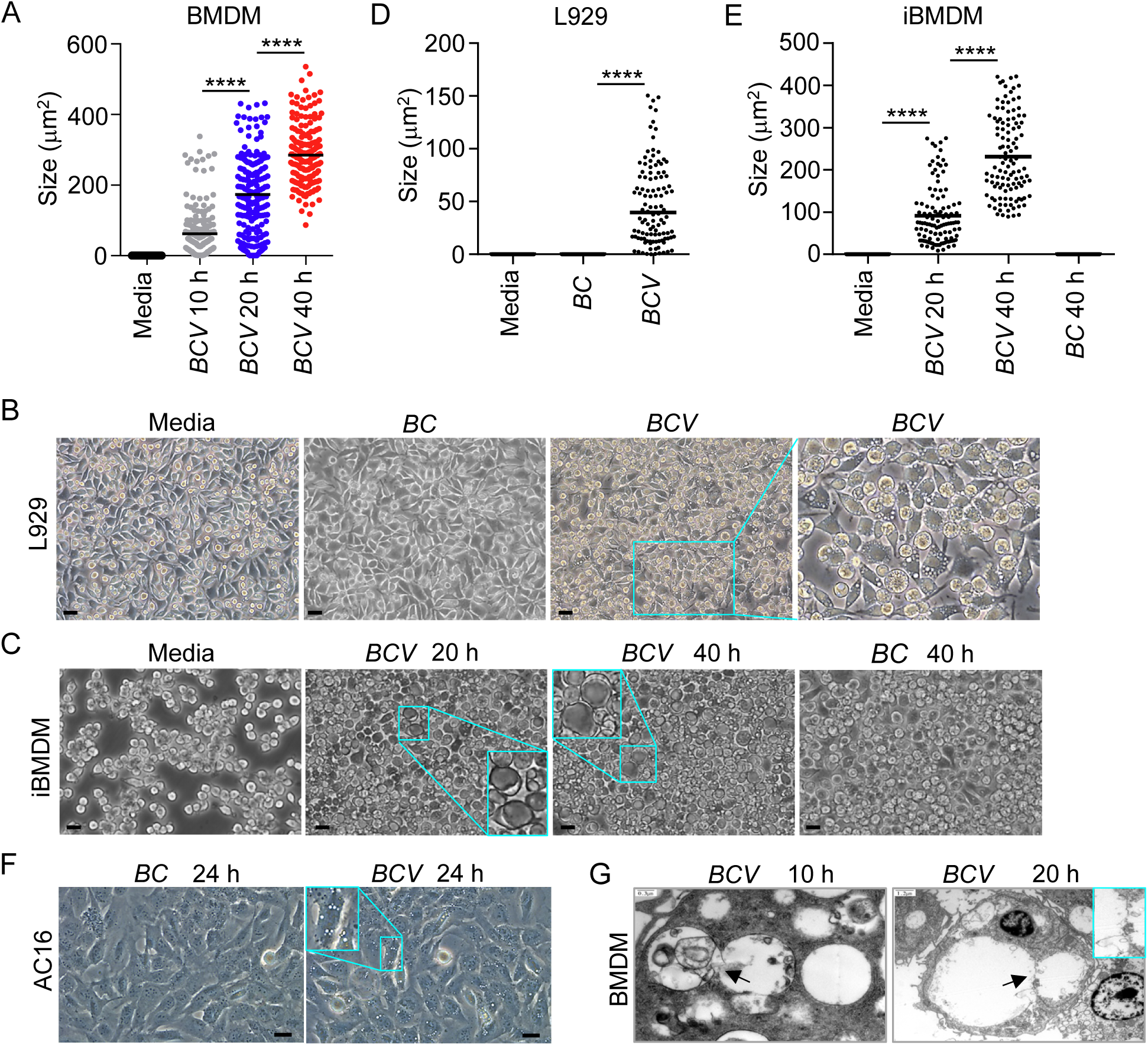
*BCV* infection induces cytoplasmic vacuolization in multiple types of cells. **A,** Quantification of vacuole size in WT BMDMs infected with *BCV* (400 MOI) for indicated times in (Figure 2A) using ImageJ. The largest vacuole per cell was analyzed, and 200 cells were quantified for each group. **B**, Microscopy analysis of L929 cells infected with *BC* or *BCV* (400 MOI) for 12 hours. Scale bars, 20 μm. **C**, Microscopy analysis of iBMDMs infected with *BC* or *BCV* (400 MOI) for the indicated times. Scale bars, 20 μm. **D**, Quantification of vacuole size in L929 cells in (**B**). The largest vacuole per cell was analyzed, and at least 110 cells were quantified for each group. **E**, Quantification of vacuole size in iBMDMs in (**C**). The largest vacuole per cell was analyzed, and at least 110 cells were quantified for each group. **F**, Microscopy analysis of AC16 infected with *BC* or *BCV* (400 MOI) for 24 hours. Scale bars, 20 μm. **G**, TEM analysis of WT BMDMs infected with *BCV* (400 MOI) for the indicated times. The arrows indicate the membrane fusion, and enlarged image is shown. Data are representative of 3 independent experiments with similar results (**A**-**G**). Data represent Mean ± SEM for (**A**, **D**, **E**), ****P < 0.0001, by 2-sided Student’s t test without multiple-comparisons correction.

**Figure S5.**
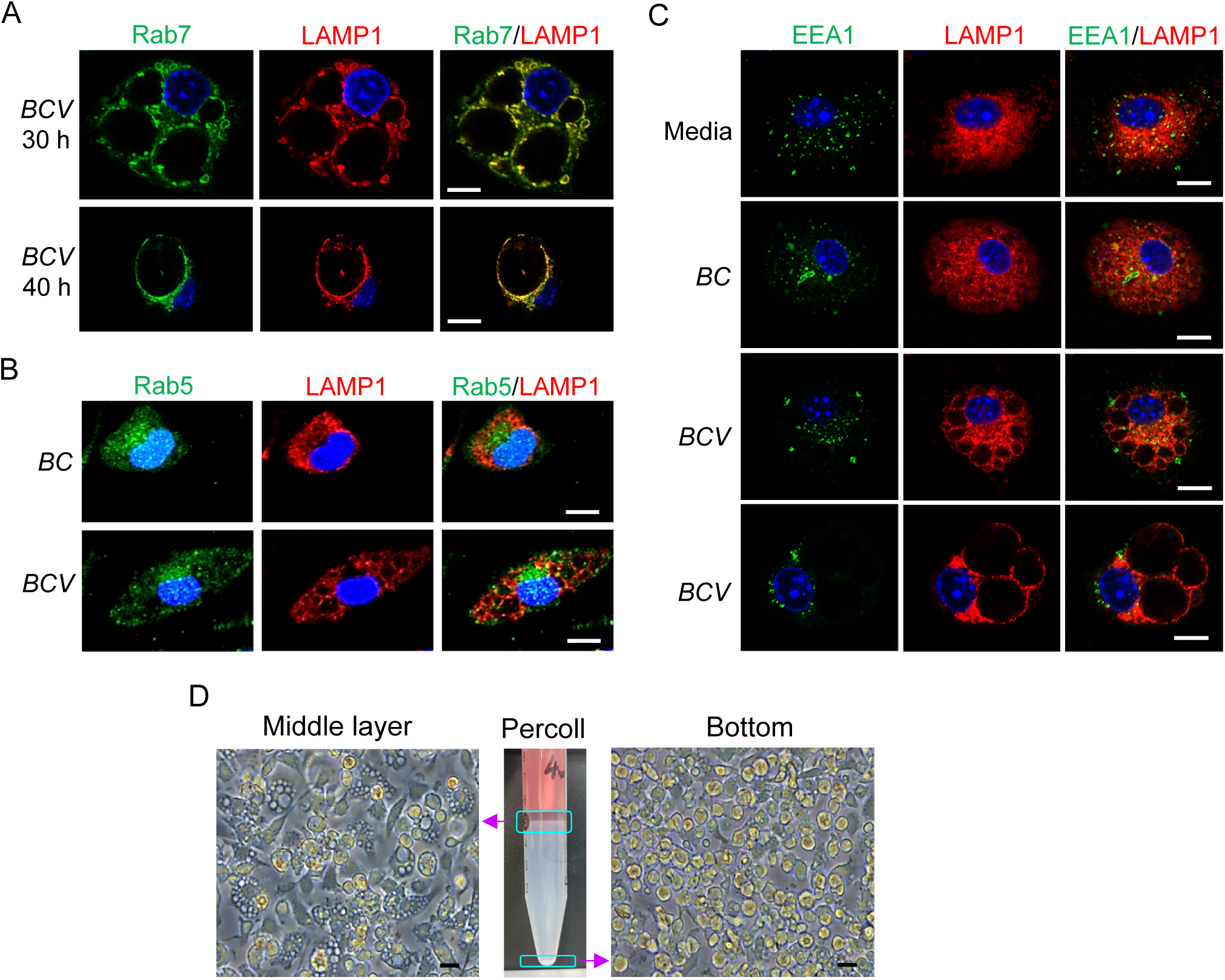
Characterization of *BCV* infection induced cell death. **A**, Confocal microscopy analysis of Rab7 and LAMP1 in *BC*- or *BCV*-infected (400 MOI) WT BMDMs for the indicated times. Scale bars, 10 μm. **B**, Confocal microscopy analysis of Rab5 and LAMP1 in *BC*- or *BCV*-infected (400 MOI) WT BMDMs for 20 hours. Scale bars, 10 μm. **C**, Confocal microscopy analysis of EEA1 and LAMP1 in uninfected (Media), and *BC*- or *BCV*-infected (400 MOI) WT BMDMs for 20 hours. Scale bars, 10 μm. **D**, Microscopy analysis of the mixture of uninfected and *BCV*-infected BMDMs (400 MOI, 20 h) separated by 40% Percoll gradient. Scale bars, 20 μm. Data are representative of 3 independent experiments with similar results (**A**-**D**).

**Figure S6.**
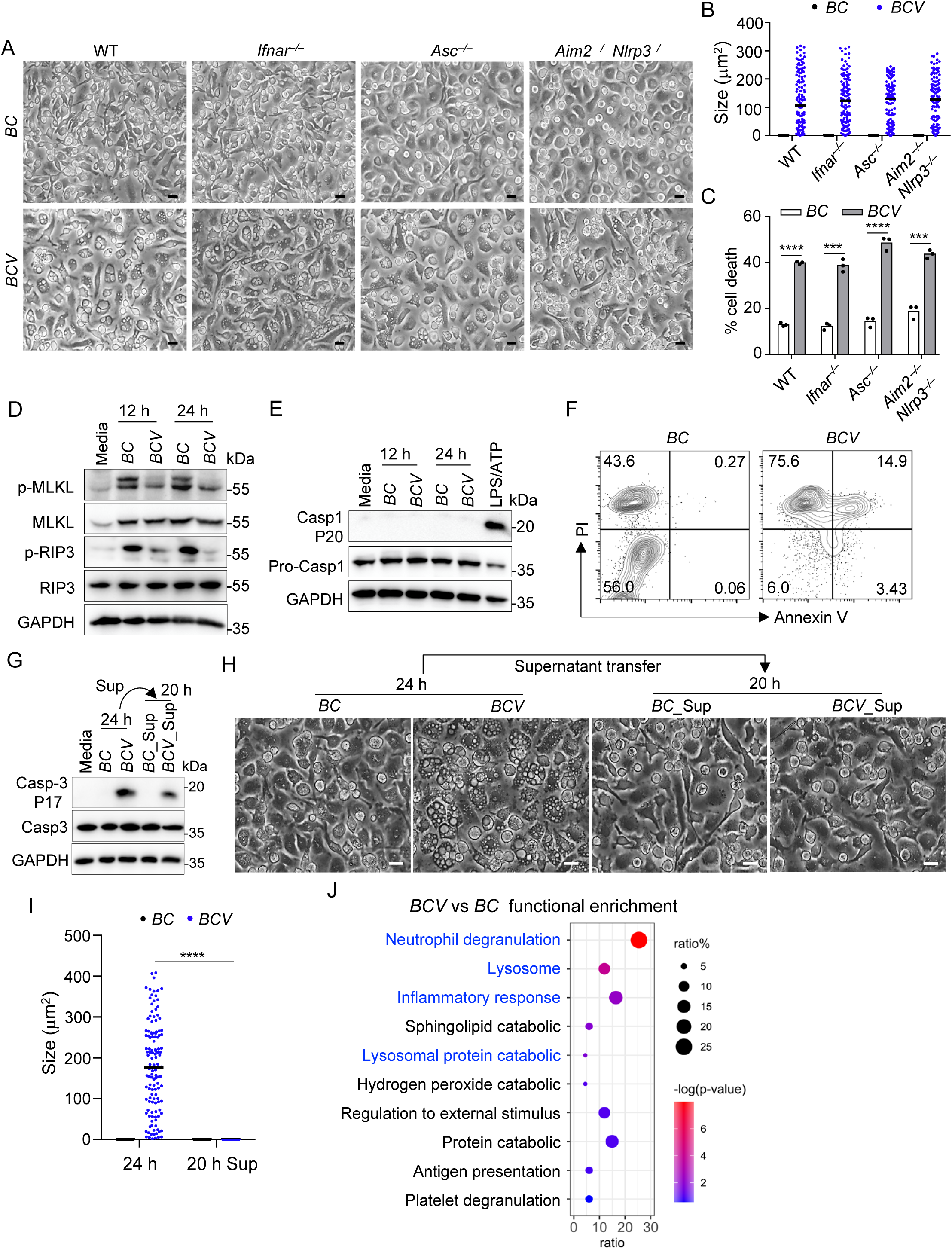
IFN-I and inflammasome are dispensable for *BCV* infection-triggered cytoplasmic vacuolization cell death. **A**, Microscopy analysis of WT, *Ifnar^-/-^, Asc^-/-^, and Aim2^-/-^Nlrp3^-/-^* BMDMs infected with *BC* or *BCV* (400 MOI) for 12 hours. Scale bars, 20 μm. **B**, Quantification of vacuole size in BMDMs in (**A**). The largest vacuole per cell was analyzed, and at least 130 cells were quantified for each group. **C**, LDH analysis of WT, *Ifnar^-/-^, Asc^-/-^, and Aim2^-/-^Nlrp3^-/-^* BMDMs infected with *BC* or *BCV* (400 MOI) for 12 hours (n=3 biologically independent samples). **D**, Immunoblot analysis of p-MLKL, MLKL, p-RIP3, and RIP3 in WT BMDMs infected with *BC* or *BCV* (400 MOI) for the indicated times. **E**, Immunoblot analysis of caspase-1 and cleaved caspase-1 (P20) in WT BMDMs infected with *BC* or *BCV* (400 MOI) for the indicated times. LPS and ATP treatment as a positive control to activate the NLRP3 inflammasome. **F**, Flow cytometry analysis of PI (10 μg/ml) and Annexin V (2 μg/ml) staining for apoptosis in WT BMDMs infected with *BC* or *BCV* (400 MOI) for 20 hours. **G**, Immunoblot analysis of caspase-3 and cleaved caspase-3 (P17) in WT BMDMs infected with *BC* or *BCV* (400 MOI) for 24 hours or treated with the supernatants (Sup) from *BC*- and *BCV*-infected BMDMs for 20 hours. Supernatants were passed through 0.22 μm filer and centrifugation to remove viable bacteria and debris. **H**, Microscopy analysis of WT BMDMs infected with *BC* or *BCV* (400 MOI) for 24 hours, or treated with supernatants from *BC*- or *BCV*-infected BMDM for 20 hours. Scale bars, 20 μm. **I**, Quantification of vacuole size in BMDMs in (**H**). The largest vacuole per cell was analyzed, and at least 130 cells were quantified for each group. **J**, Enrichment analysis of host proteins in supernatants from *BCV*- or *BC*-infected (400 MOI) BMDMs for 20 hours detected by MS. Data are from 3 independent experiments (**C**) or representative of 3 independent experiments with similar results (**A**, **B**, **D**-**I**). Data represent Mean ± SEM for (**B**, **C**, **I**), ***P < 0.001, ****P < 0.0001, by 2-sided Student’s t test without multiple- comparisons correction.

**Figure S7.**
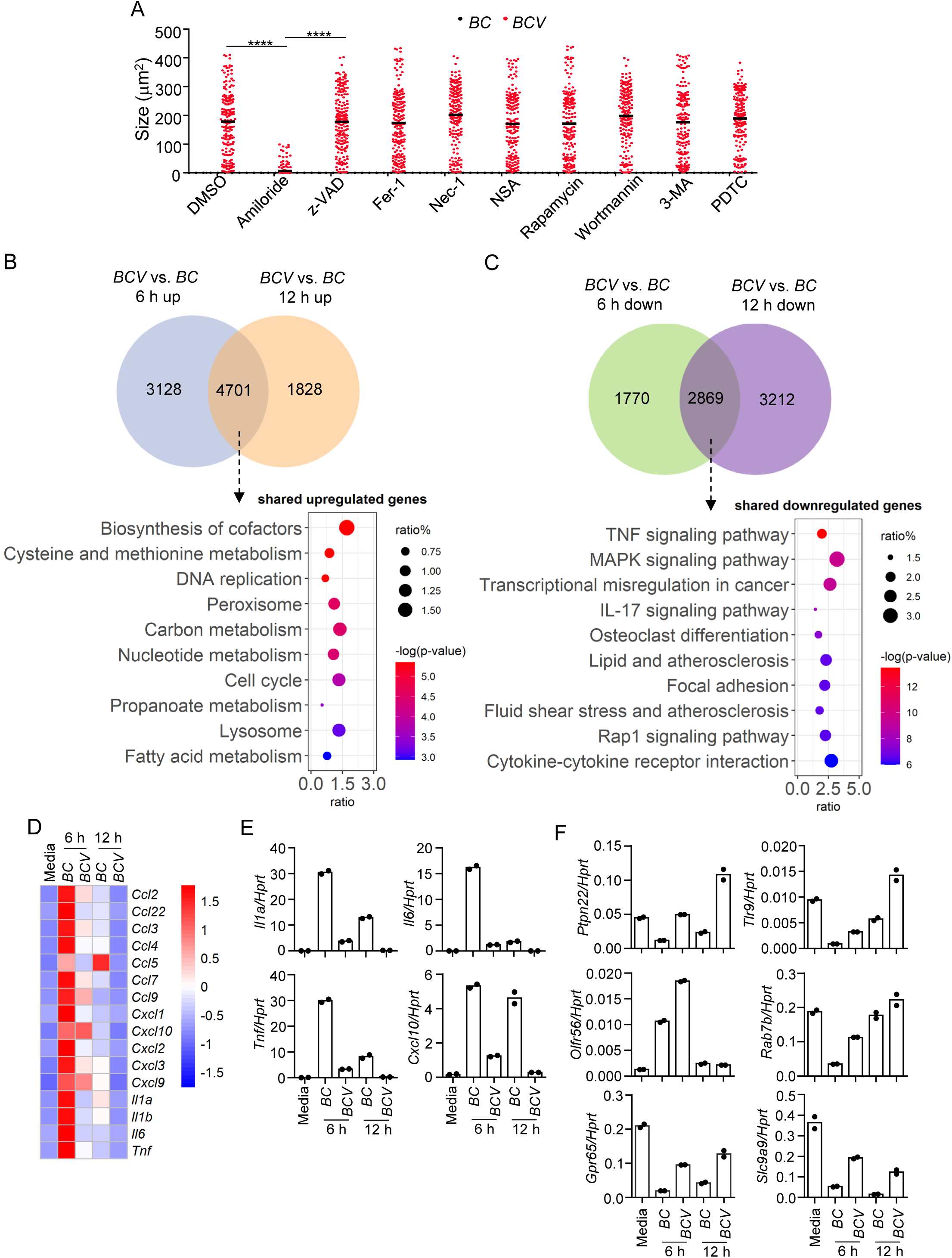
Inflammatory response is downregulated in *BCV*-infected BMDMs compared with *BC*-infected BMDMs. **A**, Quantification of vacuole size in BMDMs in (Figure 3C). The largest vacuole per cell was analyzed, and at least 200 cells were quantified for each group. **B**, Enrichment analysis of the metabolic pathways highly expressed in *BCV*-infected BMDMs. The top 10 enriched pathways of upregulated DEGs (*BCV* vs *BC*) are displayed. **C**, Enrichment analysis of the signaling pathways highly expressed in *BC*-infected BMDMs. The top 10 enriched pathways of downregulated DEGs (*BCV* vs *BC*) are displayed. **D**, Heatmap analysis of genes that encoding cytokines and chemokines with decreased expression in *BCV*-infected BMDMs. **E**, Quantitative RT-PCR analysis of *Il1a*, *Il6*, *Tnf*, and *Cxcl10* expression in uninfected (Media), and *BC*- and *BCV*-infected BMDMs for indicated times (n=2 technical replicates; 3 independent experiments). **F**, Quantitative RT-PCR analysis of *Ptpn22*, *Tlr9*, *Olfr56*, *Rab7b*, *Gpr65*, and *Slc9a9* expression in uninfected (Media), and *BC*- or *BCV*-infected BMDMs for indicated times (n=2 technical replicates; 3 independent experiments). Data are representative of 3 independent experiments with similar results (**A**, **E**, **F**). Data represent Mean ± SEM for (**A**), ****P < 0.0001, by 2-sided Student’s t test without multiple-comparisons correction.

**Figure S8.**
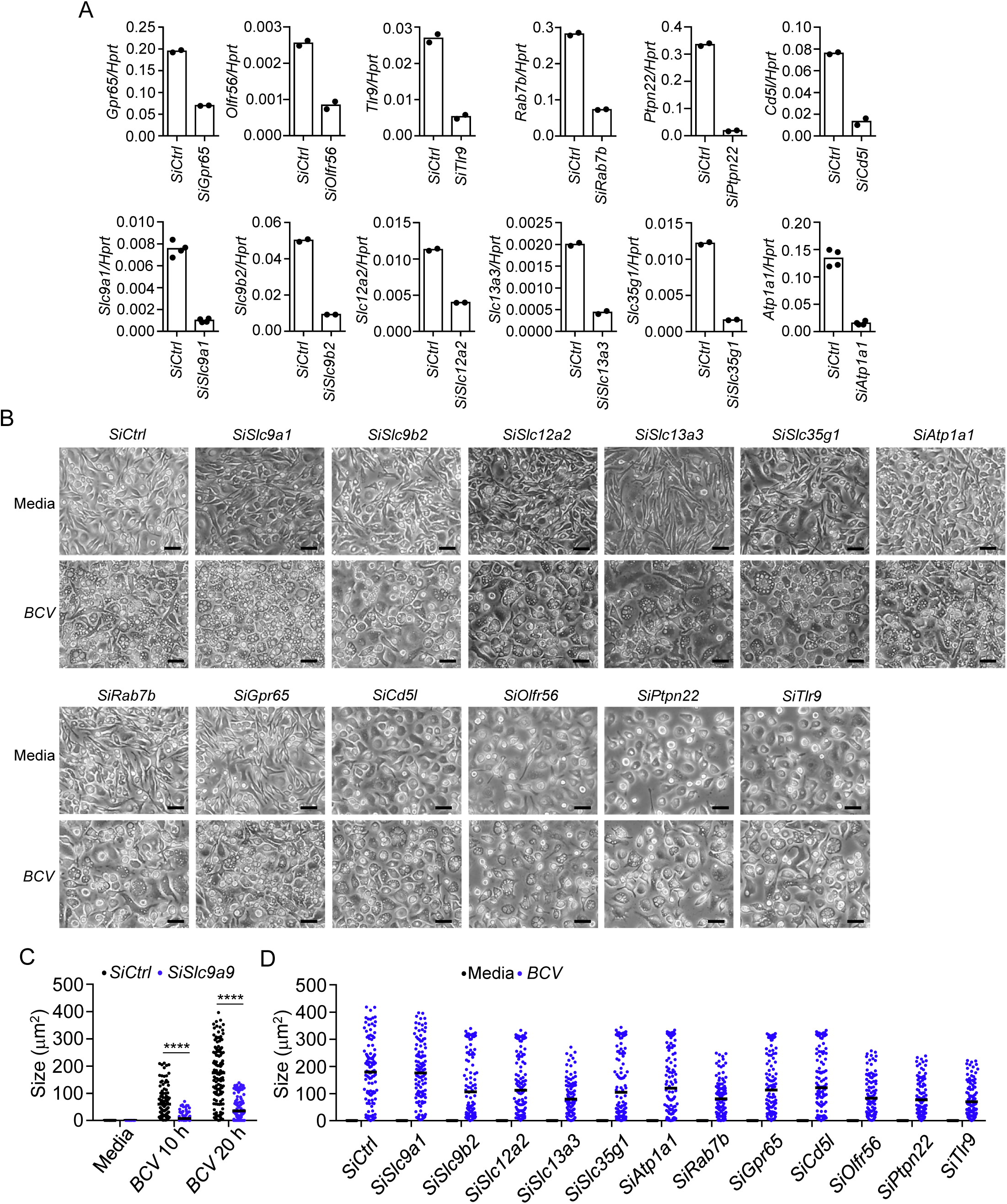
The effect of siRNAs knockdown for *BCV* infection-induced cytoplasmic vacuolization in BMDMs. **A**, Quantitative RT-PCR analysis of *Gpr65*, *Olfr56*, *Tlr9*, *Rab7b*, *Ptpn22*, *Cd5l*, *Slc9a1*, *Slc9b2*, *Slc12a2*, *Slc13a3*, *Slc35g1*, and *Atp1a1* expression in *siRNAs* transfected BMDMs (20 μM siRNA for each) for 20 hours as indicated (n=2 or 4 technical replicates; 3 independent experiments). **B**, Microscopy analysis of siRNAs-knockdown BMDMs infected with *BCV* (400 MOI) for 20 hours. Scale bars, 30 μm. **C**, Quantification of vacuole size in BMDMs in (Figure 4D). The largest vacuole per cell was analyzed, and at least 130 cells were quantified for each group. **D**, Quantification of vacuole size in BMDMs in (**B**). The largest vacuole per cell was analyzed, and at least 130 cells were quantified for each group. Data are representative of 3 independent experiments with similar results (**A**-**D**). Data represent Mean ± SEM for (**C**, **D**), ****P < 0.0001, by 2-sided Student’s t test without multiple-comparisons correction.

**Figure S9.**
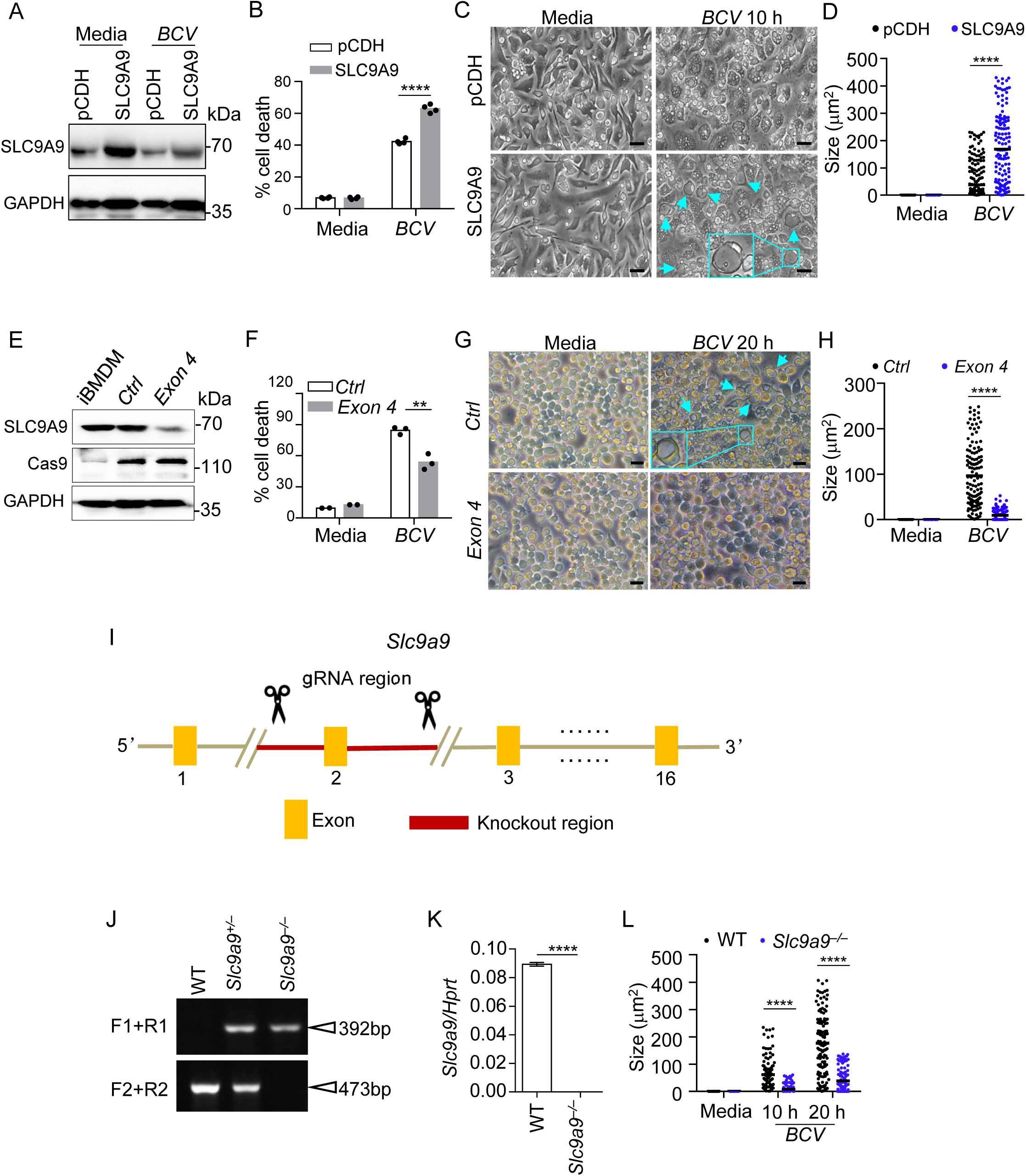
The effect of SLC9A9 for *BCV* infection-induced cytoplasmic vacuolization. **A**, Immunoblot analysis of SLC9A9 in plasmid control-(PCDH) and SLC9A9-transfected BMDMs with and without *BCV* infection (400 MOI) for 10 hours. **B**,**C**, LDH (**B**) and microscopy analysis (**C**) of plasmid control-(PCDH) and SLC9A9-transfected BMDMs with and without *BCV* infection (400 MOI) for 10 hours (n=4 biologically independent samples). The arrows indicate vacuole-occupied cells. Scale bars, 30 μm. **D**, Quantification of vacuole size in BMDMs in (**c**). The largest vacuole per cell was analyzed, and at least 130 cells were quantified for each group. **E**, Immunoblot analysis of SLC9A9 in normal iBMDMs, *Slc9a9* gene knockout (*Exon4*), control siRNA-transfected (*Ctrl*) iBMDMs by CRISPR/Cas9 transfection. **F**,**G**, LDH (**F**) and microscopy analysis (**G**) of CRISPR/Cas9-mediated *Slc9a9* gene knockout and control iBMDMs with and without *BCV* infection (400 MOI) for 20 hours (n=3 biologically independent samples). The arrows indicate vacuole-occupied cells. Scale bars, 30 μm. **H**, Quantification of vacuole size in iBMDMs in (**G**). The largest vacuole per cell was analyzed, and at least 130 cells were quantified for each group. **I**-**K**, Targeting strategy used to generate *Slc9a9^-/-^* mice (**I**), genotyping of offspring generated from breeding of *Slc9a9* heterozygous mice (**J**), and quantitative RT-PCR analysis of *Slc9a9* in WT and *Slc9a9^-/-^* BMDMs (**K**) (n=2 technical replicates; 3 independent experiments). **L**, Quantification of vacuole size in BMDMs in (Figure 4F). The largest vacuole per cell was analyzed, and at least 130 cells were quantified for each group. Data are from 3 independent experiments (**B**, **F**) or representative of 3 independent experiments with similar results (**A**, **C**, **D**, **E**, **G**, **H**, **J**-**L**). Data represent Mean ± SEM for (**B**, **D**, **F**, **H**, **K**, **L**), **P < 0.01, ****P < 0.0001, by 2-sided Student’s t test without multiple-comparisons correction.

**Figure S10.**
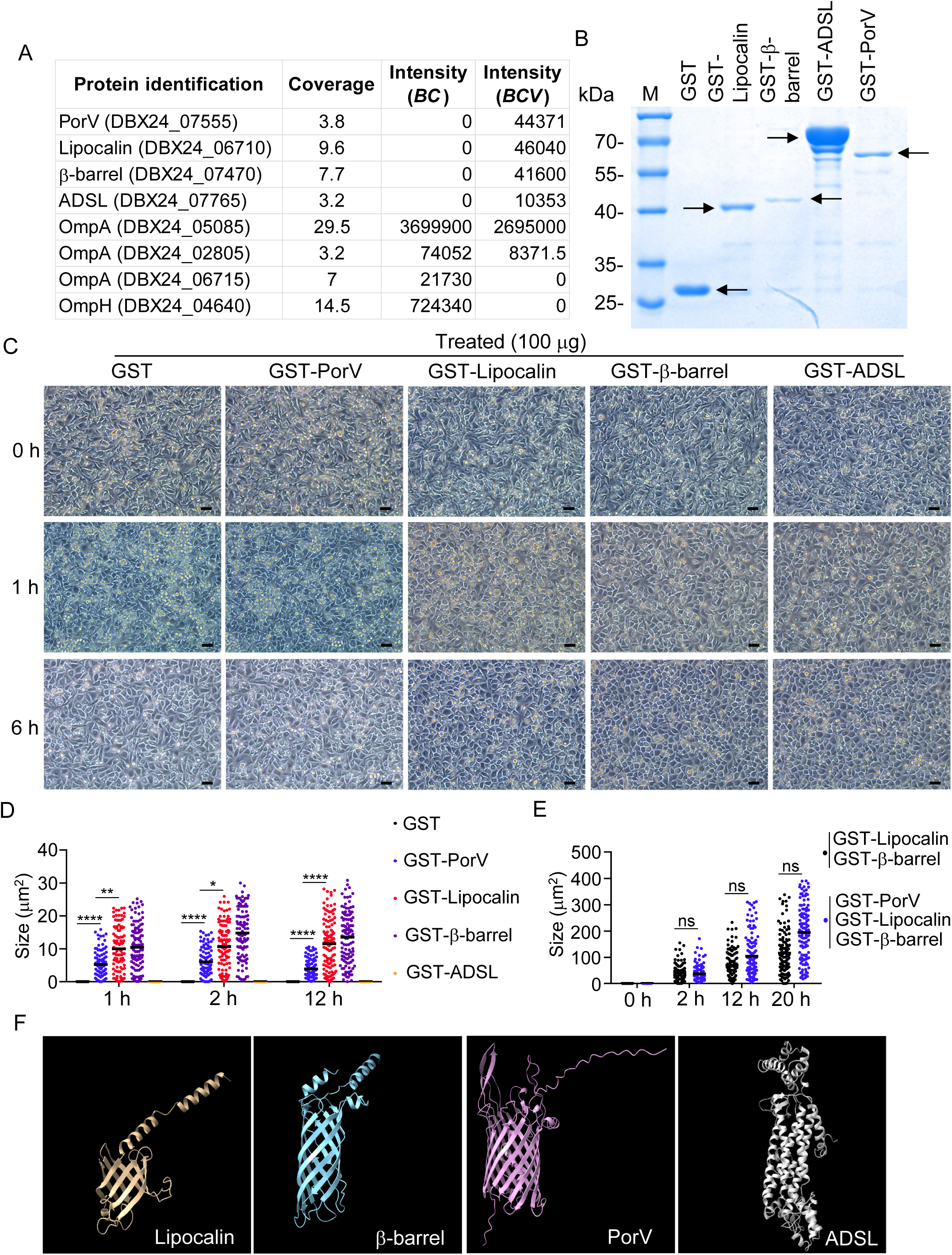
Identification and recombinant protein expression of Lipocalin, β-barrel, PorV, and ADSL from *BCV*. **A**, MS analysis of bacterial proteins in the supernatants from *BCV*- or *BC*-infected (400 MOI) BMDMs for 20 hours. **B**, SDS-PAGE of purified recombinant GST-Lipocalin, GST-β-barrel, GST-ADSL, GST-PorV, and GST control proteins, with detection by staining with coomassie blue. The arrows indicate the purified protein. **C**, Microscopy analysis of WT BMDMs treated with proteins of GST (100 μg), GST-PorV (100 μg), GST-Lipocalin (100 μg), GST-β-barrel (100 μg), and GST-ADSL (100 μg) for the indicated times. Scale bars, 30 μm. **D**, Quantification of vacuole size in BMDMs in (Figure 6A). The largest vacuole per cell was analyzed, and at least 120 cells were quantified for each group. **E**, Quantification of vacuole size in BMDMs in (Figure 6C). The largest vacuole per cell was analyzed, and at least 120 cells were quantified for each group. **F**, Predicted protein structure of Lipocalin, β-barrel, PorV, and ADSL from *BCV* by trRosseta. Data are representative of 3 independent experiments with similar results (**B**-**E**). Data represent Mean ± SEM for (**D**, **E**), *P < 0.05, **P < 0.01, ****P < 0.0001, by 2-sided Student’s t test without multiple-comparisons correction.

**Figure S11.**
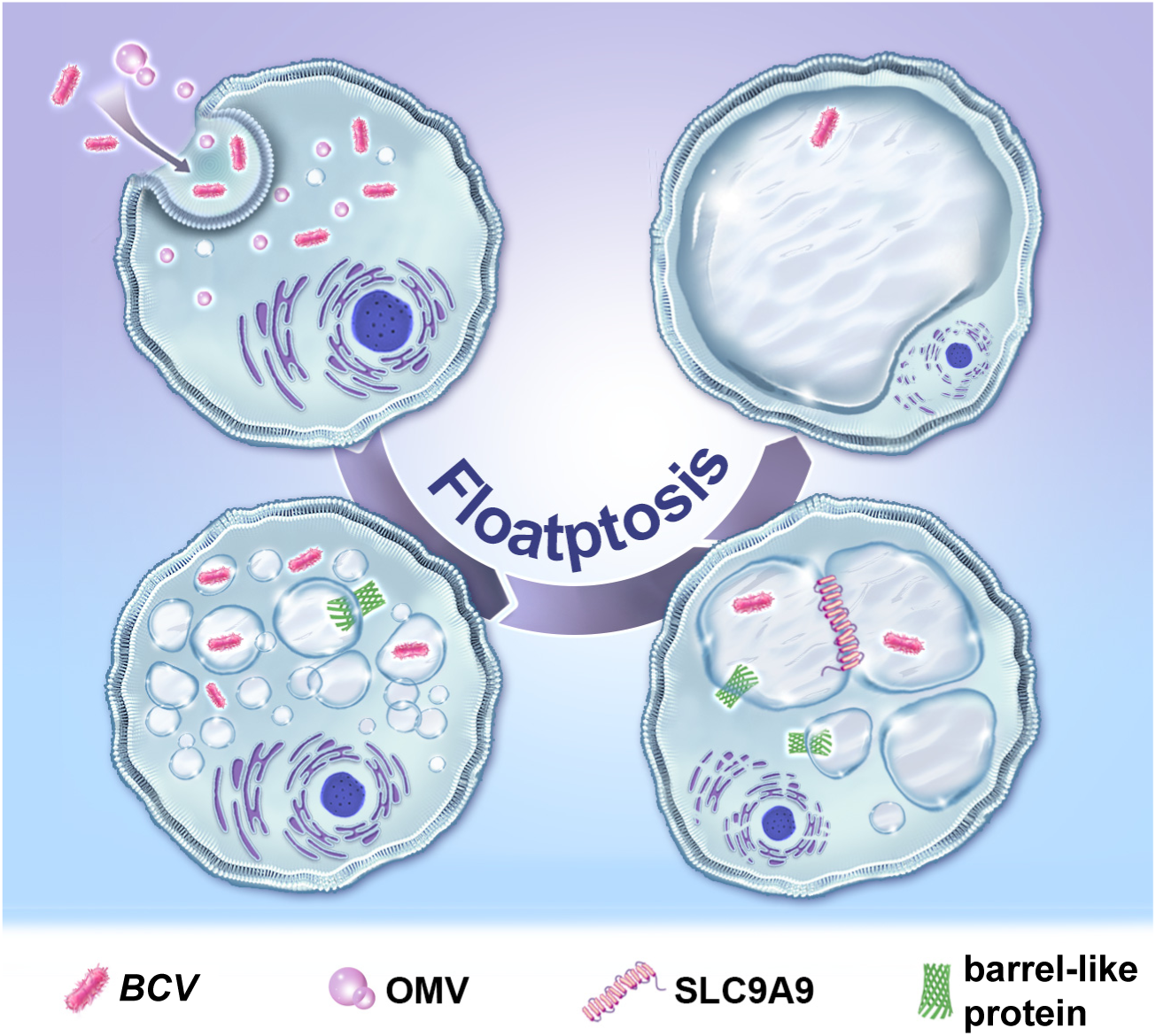
A hypothetical model of the mechanisms driving *BCV*-driven floatptosis. Bacterial pathogens have evolved multiple mechanisms to modulate host cell death. *Bergeyella cardium variant* (*BCV*) triggers unique cytoplasmic vacuolization cell death–fused lysosome-associated termination (floatptosis). The host transmembrane protein SLC9A9 and, *BCV* outer membrane vesicles (OMVs) and barrel-like proteins play important roles in the process of *BCV*-induced floatptosis.

## REFERENCES

1. Ashida, H., Mimuro, H., Ogawa, M., Kobayashi, T., Sanada, T., Kim, M., and Sasakawa, C. (2011). Cell death and infection: a double-edged sword for host and pathogen survival. J Cell Biol 195, 931–942.

2. Maudet, C., Kheloufi, M., Levallois, S., Gaillard, J., Huang, L., Gaultier, C., Tsai, Y.H., Disson, O., and Lecuit, M. (2022). Bacterial inhibition of Fas-mediated killing promotes neuroinvasion and persistence. Nature 603, 900–906.

3. Jorgensen, I., Rayamajhi, M., and Miao, E.A. (2017). Programmed cell death as a defence against infection. Nat Rev Immunol 17, 151–164.

4. Place, D.E., Lee, S., and Kanneganti, T.D. (2021). PANoptosis in microbial infection. Curr Opin Microbiol 59, 42–49.

5. Wanford, J.J., Hachani, A., and Odendall, C. (2022). Reprogramming of Cell Death Pathways by Bacterial Effectors as a Widespread Virulence Strategy. Infect Immun 90, e0061421.

6. Shubin, A.V., Demidyuk, I.V., Komissarov, A.A., Rafieva, L.M., and Kostrov, S.V. (2016). Cytoplasmic vacuolization in cell death and survival. Oncotarget 7, 55863–55889.

7. Pan, H., Li, W., Sun, E., and Zhang, Y. (2020). Characterization and whole genome sequencing of a novel strain of Bergeyella cardium related to infective endocarditis. BMC microbiology 20, 32.

8. Ledger, E.V.K., Mesnage, S., and Edwards, A.M. (2022). Human serum triggers antibiotic tolerance in Staphylococcus aureus. Nature Communications 13, 2041.

9. Miajlovic, H., and Smith, S.G. (2014). Bacterial self-defence: how Escherichia coli evades serum killing. FEMS Microbiology Letters 354, 1–9.

10. Weber, B.S., De Jong, A.M., Guo, A.B.Y., Dharavath, S., French, S., Fiebig-Comyn, A.A., Coombes, B.K., Magolan, J., and Brown, E.D. (2020). Genetic and Chemical Screening in Human Blood Serum Reveals Unique Antibacterial Targets and Compounds against Klebsiella pneumoniae. Cell Rep 32, 107927.

11. Lauber, F., Deme, J.C., Lea, S.M., and Berks, B.C. (2018). Type 9 secretion system structures reveal a new protein transport mechanism. Nature 564, 77–82.

12. Du, Z., Su, H., Wang, W., Ye, L., Wei, H., Peng, Z., Anishchenko, I., Baker, D., and Yang, J. (2021). The trRosetta server for fast and accurate protein structure prediction. Nature Protocols 16, 5634–5651.

13. Guo, Y., Li, L., Xu, T., Guo, X., Wang, C., Li, Y., Yang, Y., Yang, D., Sun, B., Zhao, X., et al. (2020). HUWE1 mediates inflammasome activation and promotes host defense against bacterial infection. The Journal of Clinical Investigation 130, 6301–6316.

14. Li, Y., Guo, X., Hu, C., Du, Y., Guo, C., Di, W., Zhao, W., Huang, G., Li, C., Lu, Q., et al. (2018). Type I IFN operates pyroptosis and necroptosis during multidrug-resistant A. baumannii infection. Cell Death Differ 25, 1304–1318.

15. Mulliken, J.S., Langelier, C., Budak, J.Z., Miller, S., Dynerman, D., Hao, S., Li, L.M., Crawford, E., Lyden, A., Woodworth, M.H., et al. (2019). Bergeyella cardium: Clinical Characteristics and Draft Genome of an Emerging Pathogen in Native and Prosthetic Valve Endocarditis. Open forum infectious diseases 6, ofz134.

16. Kaparakis-Liaskos, M., and Ferrero, R.L. (2015). Immune modulation by bacterial outer membrane vesicles. Nature Reviews Immunology 15, 375–387.

17. Devireddy, L.R., Teodoro, J.G., Richard, F.A., and Green, M.R. (2001). Induction of Apoptosis by a Secreted Lipocalin That is Transcriptionally Regulated by IL-3 Deprivation. Science 293, 829–834.

18. Noinaj, N., Kuszak, A.J., Gumbart, J.C., Lukacik, P., Chang, H., Easley, N.C., Lithgow, T., and Buchanan, S.K. (2013). Structural insight into the biogenesis of β-barrel membrane proteins. Nature 501, 385–390.

19. Vorobieva, A.A., White, P., Liang, B., Horne, J.E., Bera, A.K., Chow, C.M., Gerben, S., Marx, S., Kang, A., Stiving, A.Q., et al. (2021). De novo design of transmembrane β barrels. Science 371.

20. Mandal, K. (2020). Review of PIP2 in Cellular Signaling, Functions and Diseases. Int J Mol Sci 21.

21. Zolov, S.N., Bridges, D., Zhang, Y., Lee, W.-W., Riehle, E., Verma, R., Lenk, G.M., Converso-Baran, K., Weide, T., Albin, R.L., et al. (2012). In vivo, Pikfyve generates PI(3,5)P2, which serves as both a signaling lipid and the major precursor for PI5P. Proceedings of the National Academy of Sciences 109, 17472–17477.

22. Chow, C.Y., Zhang, Y., Dowling, J.J., Jin, N., Adamska, M., Shiga, K., Szigeti, K., Shy, M.E., Li, J., Zhang, X., et al. (2007). Mutation of FIG4 causes neurodegeneration in the pale tremor mouse and patients with CMT4J. Nature 448, 68–72. 10.1038/nature05876.

23. Lenk, G.M., and Meisler, M.H. (2014). Mouse models of PI(3,5)P2 deficiency with impaired lysosome function. Methods in enzymology 534, 245–260. 10.1016/b978-0-12-397926-1.00014-7.

24. Lees, J.A., Li, P., Kumar, N., Weisman, L.S., and Reinisch, K.M. (2020). Insights into Lysosomal PI(3,5)P(2) Homeostasis from a Structural-Biochemical Analysis of the PIKfyve Lipid Kinase Complex. Mol Cell 80, 736–743.e734. 10.1016/j.molcel.2020.10.003.

25. Lenk, G.M., Szymanska, K., Debska-Vielhaber, G., Rydzanicz, M., Walczak, A., Bekiesinska-Figatowska, M., Vielhaber, S., Hallmann, K., Stawinski, P., Buehring, S., et al. (2016). Biallelic Mutations of VAC14 in Pediatric-Onset Neurological Disease. American journal of human genetics 99, 188–194. 10.1016/j.ajhg.2016.05.008.

26. Campeau, Philippe M., Lenk, Guy M., Lu, James T., Bae, Y., Burrage, L., Turnpenny, P., Román Corona-Rivera, J., Morandi, L., Mora, M., Reutter, H., et al. (2013). Yunis-Varon Syndrome Is Caused by Mutations in FIG4, Encoding a Phosphoinositide Phosphatase. The American Journal of Human Genetics 92, 781–791. 10.1016/j.ajhg.2013.03.020.

27. Zhang, Y., Zolov, S.N., Chow, C.Y., Slutsky, S.G., Richardson, S.C., Piper, R.C., Yang, B., Nau, J.J., Westrick, R.J., Morrison, S.J., et al. (2007). Loss of Vac14, a regulator of the signaling lipid phosphatidylinositol 3,5-bisphosphate, results in neurodegeneration in mice. Proc Natl Acad Sci U S A 104, 17518–17523. 10.1073/pnas.0702275104.

28. Stewart, P.E., Carroll, J.A., Dorward, D.W., Stone, H.H., Sarkar, A., Picardeau, M., and Rosa, P.A. (2012). Characterization of the Bat proteins in the oxidative stress response of Leptospira biflexa. BMC microbiology 12, 290.

29. Dieppedale, J., Sobral, D., Dupuis, M., Dubail, I., Klimentova, J., Stulik, J., Postic, G., Frapy, E., Meibom, K.L., Barel, M., and Charbit, A. (2011). Identification of a putative chaperone involved in stress resistance and virulence in Francisella tularensis. Infect Immun 79, 1428–1439.

30. Veith, P.D., Glew, M.D., Gorasia, D.G., and Reynolds, E.C. (2017). Type IX secretion: the generation of bacterial cell surface coatings involved in virulence, gliding motility and the degradation of complex biopolymers. Molecular Microbiology 106, 35–53.

31. Veith, P.D., Glew, M.D., Gorasia, D.G., Cascales, E., and Reynolds, E.C. (2022). The Type IX Secretion System and Its Role in Bacterial Function and Pathogenesis. Journal of Dental Research 101, 374–383.

32. Kim, H.-M., and Davey, M.E. (2020). Synthesis of ppGpp impacts type IX secretion and biofilm matrix formation in Porphyromonas gingivalis. npj Biofilms and Microbiomes 6, 5.

33. Sheridan, P.O., Odat, M.A., and Scott, K.P. (2023). Establishing genetic manipulation for novel strains of human gut bacteria. Microbiome research reports 2, 1. 10.20517/mrr.2022.13.

34. Morissette, G., Moreau, E., R, C.G., and Marceau, F. (2004). Massive cell vacuolization induced by organic amines such as procainamide. The Journal of pharmacology and experimental therapeutics 310, 395–406.

35. Marceau, F., Bawolak, M.-T., Lodge, R., Bouthillier, J., Gagné-Henley, A., C.- Gaudreault, R., and Morissette, G. (2012). Cation trapping by cellular acidic compartments: Beyond the concept of lysosomotropic drugs. Toxicology and Applied Pharmacology 259, 1–12.

36. Liu, Y., Shoji-Kawata, S., Sumpter, R.M., Jr., Wei, Y., Ginet, V., Zhang, L., Posner, B., Tran, K.A., Green, D.R., Xavier, R.J., et al. (2013). Autosis is a Na+,K+-ATPase-regulated form of cell death triggered by autophagy-inducing peptides, starvation, and hypoxia-ischemia. Proc Natl Acad Sci U S A 110, 20364–20371.

37. Fuster, D.G., and Alexander, R.T. (2014). Traditional and emerging roles for the SLC9 Na+/H+ exchangers. Pflügers Archiv - European Journal of Physiology 466, 61–76.

38. Zhang-James, Y., Vaudel, M., Mjaavatten, O., Berven, F.S., Haavik, J., and Faraone, S.V. (2019). Effect of disease-associated SLC9A9 mutations on protein-protein interaction networks: implications for molecular mechanisms for ADHD and autism. Attention deficit and hyperactivity disorders 11, 91–105.

39. Winkelmann, I., Matsuoka, R., Meier, P.F., Shutin, D., Zhang, C., Orellana, L., Sexton, R., Landreh, M., Robinson, C.V., Beckstein, O., and Drew, D. (2020). Structure and elevator mechanism of the mammalian sodium/proton exchanger NHE9. The EMBO Journal 39, e105908.

40. Chen, R., and Chen, L. (2022). Solute carrier transporters: emerging central players in tumour immunotherapy. Trends Cell Biol 32, 186–201.

41. Song, W., Li, D., Tao, L., Luo, Q., and Chen, L. (2020). Solute carrier transporters: the metabolic gatekeepers of immune cells. Acta pharmaceutica Sinica. B 10, 61–78.

42. Xu, H., Ghishan, F.K., and Kiela, P.R. (2018). SLC9 Gene Family: Function, Expression, and Regulation. Comprehensive Physiology 8, 555–583.

43. Pedersen, S.F., and Counillon, L. (2019). The SLC9A-C Mammalian Na(+)/H(+) Exchanger Family: Molecules, Mechanisms, and Physiology. Physiol Rev 99, 2015–2113.

44. Kondapalli, K.C., Hack, A., Schushan, M., Landau, M., Ben-Tal, N., and Rao, R. (2013). Functional evaluation of autism-associated mutations in NHE9. Nat Commun 4, 2510.

45. Gurnari, C., Pagliuca, S., Durkin, L., Terkawi, L., Awada, H., Kongkiatkamon, S., Zawit, M., Hsi, E.D., Carraway, H.E., Rogers, H.J., et al. (2021). Vacuolization of hematopoietic precursors: an enigma with multiple etiologies. Blood 137, 3685–3689.

46. Guo, Y., Mao, R., Xie, Q., Cheng, X., Xu, T., Wang, X., Du, Y., and Qi, X. (2021). Francisella novicida Mutant XWK4 Triggers Robust Inflammasome Activation Favoring Infection. Front Cell Dev Biol 9:743335.

47. Chowdhury, R., Bouatta, N., Biswas, S., Floristean, C., Kharkar, A., Roy, K., Rochereau, C., Ahdritz, G., Zhang, J., Church, G.M., et al. (2022). Single-sequence protein structure prediction using a language model and deep learning. Nature Biotechnology 40, 1617–1623.

48. Su, H., Wang, W., Du, Z., Peng, Z., Gao, S.-H., Cheng, M.-M., and Yang, J. (2021). Improved Protein Structure Prediction Using a New Multi-Scale Network and Homologous Templates. Advanced Science 8, 2102592.

49. Pettersen, E.F., Goddard, T.D., Huang, C.C., Couch, G.S., Greenblatt, D.M., Meng, E.C., and Ferrin, T.E. (2004). UCSF Chimera--a visualization system for exploratory research and analysis. Journal of computational chemistry 25, 1605–1612.

